# Ultra-parallel ribosome profiling platform with RNA-dependent RNA amplification

**DOI:** 10.1101/2025.08.06.668979

**Authors:** Yuichi Shichino, Mari Mito, Yasuha Kinugasa, Taisei Wakigawa, Akira Yamashita, Yuichiro Mishima, Yumiko Imai, Shintaro Iwasaki

## Abstract

Translation regulation plays a pivotal role in the diversification of gene expression and the response to intra- and extracellular environmental cues. Ribosome profiling (or Ribo-Seq) serves as a sensitive, quantitative, comprehensive, and data-rich technique to survey ribosome traversal across the cellular transcriptome. However, due to the intricacy of library preparation, applications to low-input and a large number of samples have presented analytic challenges. Here, we developed the semi-automated platform of Ribo-Seq and Disome-Seq, which allowed us to assess the translation status from a vast collection of samples with reduced amounts in a plate format. Through an siRNA-mediated knockdown screen for ribosome-associated proteins, this technique identified factors that (i) mediate regulation via RNA elements such as the TOP motif, (ii) assist efficient ribosome recycling, (iii) support ribosome-associated quality control (RQC), (iv) enhance translation elongation across inhibitory G-quadruplex sequences, and (v) repress mitochondrial translation. The application to human-derived samples revealed that stop codon readthrough and mitochondrial translation deficiency are associated with severe symptoms in COVID-19. Our approach provides a versatile option to investigate the translatome in a highly parallel manner.

## Introduction

Although the transcriptomic profile has been recognized as a proxy for gene expression, recent studies have shown that regulation at the level of protein synthesis has a profound effect on the final output of genes. Ribosome profiling (or Ribo-Seq), a technique based on RNase footprinting of mRNAs by ribosomes and subsequent deep sequencing, has emerged as a game-changing approach for studying protein synthesis in cells ^1^. The applications of this technique are highly manifold; it can measure translation efficiencies across the transcriptome ^2,3^, assign open reading frames (ORFs) *de novo* ^4^, explore ribosome movement speed at codon resolution ^2,3^, and even estimate the structural status of ribosomes ^5,6^. The widespread applications of this technique in diverse materials and biological conditions have unveiled previously uncharacterized layers of regulation of protein synthesis ^2–4^. The road ahead for Ribo-Seq should involve the application to rare, precious, and difficult-to-prepare samples. However, standard protocols require a significant number of cells for high-quality library preparation ^7,8^ and thus present technical hurdles for low inputs.

Therefore, efforts to optimize Ribo-Seq with small amounts of materials are underway ^9–17^. A one-pot reaction for library preparation is advantageous to avoid sample loss. For this purpose, a ligation-free method could be applied to isolated ribosome footprints; this system allows the sequential reactions of 3′ end poly(A) tailing, reverse transcription, and linker addition by template switching in the same tube ^9,11–13^. However, these approaches do not have any means to amplify the DNA or RNA before PCR and thus involve the risk of material loss during the procedures.

Another hurdle in Ribo-Seq has been sample number scalability. The isolation of ribosome-footprint complexes through the sucrose cushion ultracentrifugation generally limits the sample number that can be handled in a batch (typically, ∼8 samples held in the rotor). Moreover, footprint size selection by gel purification is another labor-intensive step that restricts the sample numbers (Figure 2A). These issues pose a challenge when converting Ribo-Seq to a high-throughput format.

Here, we developed a 96-well plate-based, semi-automated, low-input tailored system of Ribo-Seq, termed high-throughput (HT) T7 High-resolution original RNA (Thor)-Ribo-Seq. This technique harnesses RNA-dependent RNA amplification by T7 RNA polymerase, which can use an RNA‒DNA chimera as an *in vitro* transcription template. Moreover, by exchanging batch-to-batch ultracentrifugation to size-exclusion gel filtration for ribosome-footprint complex purification, Thor-Ribo-Seq was formatted on 96-well plates and handled with pipetting robots. Due to the high capacity for large sample numbers and adaptability to low input, our semi-automated framework monitored translational changes by knockdown for ∼180 ribosome-binding proteins, categorizing and predicting their functions in protein synthesis. We screened translation initiation regulators through RNA motifs, elongation regulators functioning in ribosome-associated quality control (RQC), and ribosome recycling regulators. Our study identified a repressor of mitochondrial translation. Application of disome profiling in the system (*i.e.*, HT-Thor-Disome-Seq) revealed factors that suppress RNA G quadruplex-mediated ribosome collision. Expansion of our analysis to mitochondrial translation identified IGF2BP3 as a mitochondrial translational repressor. Furthermore, profiling of human peripheral blood mononuclear cells (PBMCs) derived from ∼70 COVID-19 patients revealed the symptom severity-associated regulations, such as stop codon readthrough in cytosolic translation and global inefficiency of mitochondrial translation. Our platform expands the genome-wide survey of translation to a systematic scale.

## Results and discussion

### Implementation of RNA-dependent RNA amplification into Ribo-Seq

An apparent pitfall of Ribo-Seq is the number of cells for library preparation. Indeed, standard Ribo-Seq experiments (Figure S1A) typically required 1-10 μg of total RNA, which corresponds to approximately 10^5^-10^6^ cells of human embryonic kidney (HEK) 293. Lower quantities of cell lysate, such as extract corresponding to 0.1 μg of total RNA (or ∼10^4^ cells), could not enable DNA library amplification by PCR (Figure S1C) due to sample loss during the multiple complicated steps in library generation. This drawback ultimately restricted Ribo-Seq experiments to precious, limited samples.

To overcome this issue, we developed an RNA-dependent RNA amplification strategy for ribosome footprints at an early stage of library preparation, especially before the loss of material (Figures 1A and S1B). We designed the linker DNA oligonucleotide with the antisense sequence of the T7 promoter. Hybridization of short DNA oligonucleotides covering the antisense sequence created a partial dsDNA region competent for RNA transcription ^18^. Since T7 polymerase can synthesize complementary RNA from the RNA template conjugated with the dsDNA T7 promoter region ^19,20^, this configuration allows *in vitro* transcription of the complementary RNA of ribosome footprints. To suppress the amplification bias, we added a unique molecular index (UMI) in the linkers (Figure S1B) (see below for details). We termed our strategy as T7 High-resolution Original RNA (Thor)-Ribo-Seq (inspired by the technique used in the LUTHOR 3′ mRNA-Seq Library Prep Kit provided by Lexogen).

**Figure 1.**
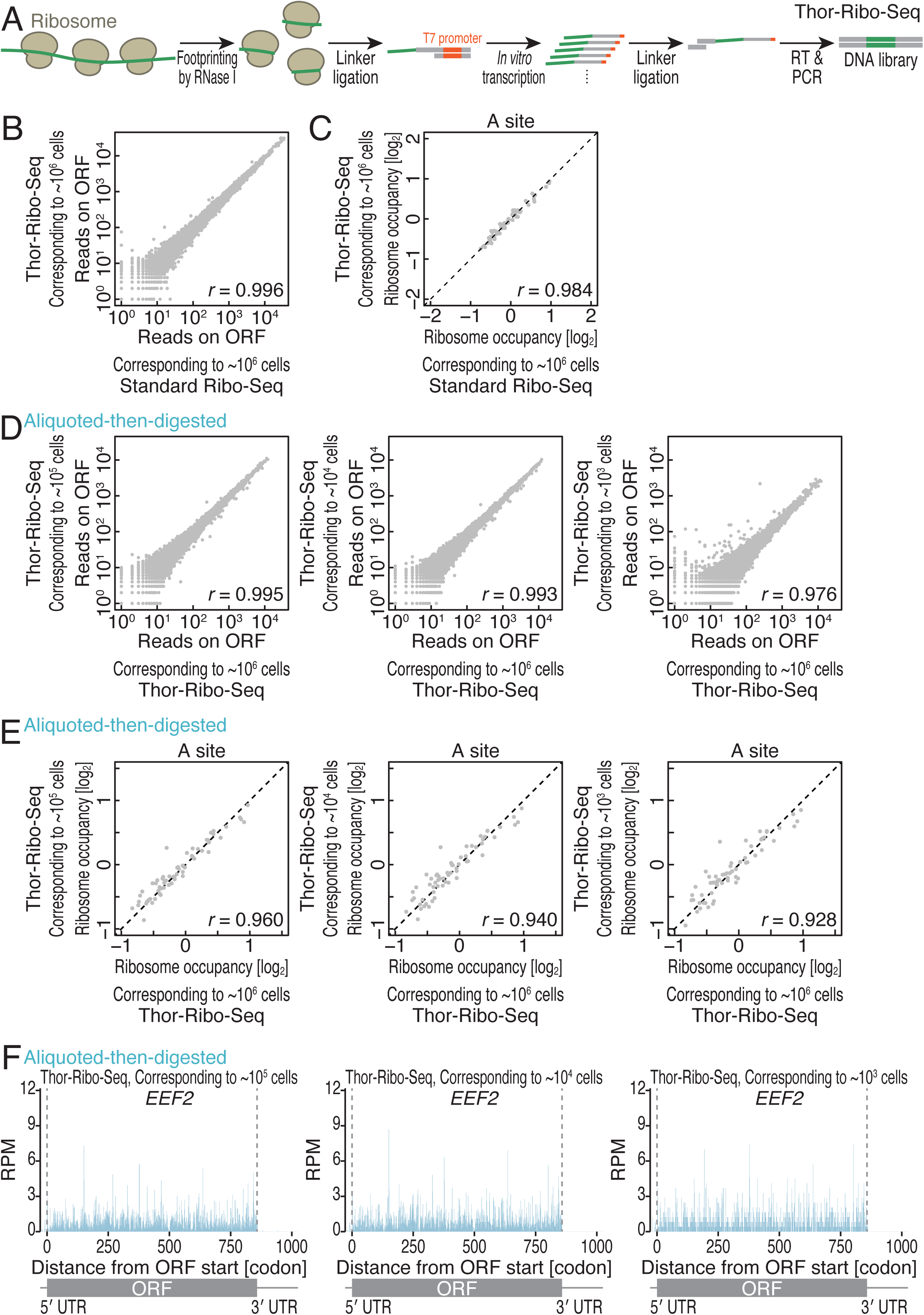
Thor-Ribo-Seq assesses translation status from low inputs. (A) Schematic representation of the library preparation strategy of Thor-Ribo-Seq. RT, reverse transcription. (B) Correspondence of reads on ORFs between standard Ribo-Seq and Thor-Ribo-Seq with a normal amount of inputs. (C) Correspondence of averaged ribosome occupancy on A-site codon sequences between standard Ribo-Seq data and Thor-Ribo-Seq with a normal amount of inputs. (D) Correspondence of reads on ORFs across Thor-Ribo-Seq with a normal amount of input and that of “aliquoted-then-digested” experiments with low inputs. (E) Correspondence of averaged ribosome occupancy on A-site codon sequences across Thor-Ribo-Seq with a normal amount of input and that of “aliquoted-then-digested” experiments with low inputs. (F) Ribosome footprint distribution on *EEF2* mRNA of Thor-Ribo-Seq with “aliquoted-then-digested” experiments with low material inputs. *r*, Pearson’s correlation coefficient. RPM, reads per million mapped reads. See also Figure S1 and Table S6.

We set out to compare the performance of the Thor-Ribo-Seq with standard Ribo-Seq with the normal amount of material (10 μg of total RNA or ∼10^6^ cells). We observed hallmarks of ribosome footprints in Thor-Ribo-Seq: read peaks at 22 and 28 nucleotides (nt) (Figure S1D) ^5,6^ and strong triplet periodicity along the ORFs(Figure S1E-F) as in standard Ribo-Seq. Also, both Ribo-Seq methods detected comparable numbers of genes (Figure S1G). Thus, high correspondence of the reads on ORFs was found between standard Ribo-Seq and Thor-Ribo-Seq (Figure 1B).

Moreover, Thor-Ribo-Seq exhibited limited artifacts in codon-wise examinations of ribosome occupancy. A-site codon sequences are the most prominent determinants of ribosome duration time at each codon ^21–30^. Thus, differential ribosome occupancies were observed (Figure 1C). This divergence was consistently maintained in Thor-Ribo-Seq (Figure 1C). The previous studies have shown that technical factors, including the sequence preference of ligase, may generate biases ^27,31^. CircLigaseII, which adds a primer landing site at the 5′ end of the footprint (Figure S1A), may prefer an “AUG” sequence at the 5′ end and thus overrepresent ribosome footprints at the 5th codons in the standard Ribo-Seq (Figure S1H). However, Thor-Ribo-Seq avoided this enzyme and harnesses second RNA-to-DNA ligation (Figure S1B) and thus suppressed such a disproportionate presentation at the 5th codons (Figure S1H).

T7 polymerase is known to extend RNA at the 3′ end ^18,32–34^, which stems from RNA self-templated extension through folding back on itself ^35^. However, the Thor protocol did not extend the footprint length (Figure S1D). This was possibly due to the tight hybridization of transcribed RNAs to template footprint RNAs, preventing further looping back.

These results indicate that the Thor-Ribo-Seq strategy provides a less-biased option.

### Thor-Ribo-Seq for translational profiling from low material input

Strikingly, the Thor-Ribo-Seq strategy allowed us to perform experiments with inputs too low to be processed by standard Ribo-Seq. Given that RNase treatment ^11,36–38^ and the downstream processes ^23,27^ introduce different biases, we designed experiments to control the two impactful sources individually when Thor-Ribo-Seq is applied to small inputs. To maintain the same RNase treatment conditions, we prepared a library with aliquots of RNase-treated lysate (denoted as “digested-then-aliquoted”) (Figure S1I); we treated cell lysate containing 10 μg of total RNA (or ∼10^6^ cells) with RNase I, as in standard Ribo-Seq, and then divided the reaction into aliquots corresponding to 0.01, 0.1, or 1 μg of total RNA for downstream Thor-Ribo-Seq library preparation. Transcription by T7 polymerase enabled the amplification of the complementary RNA (Figure S1J) and thus the sequencing library (Figure S1K), for inputs at least as low as 0.01 μg (equal to ∼10^3^ cells). The benchmarks of footprints, including read length (Figure S1L) and 3-nt periodicity (Figure S1M-O), were maintained for low-input libraries. Moreover, the reads on ORFs exhibited high correlations to the Thor-Ribo-Seq data prepared with ∼10^6^ cells (Figure S1P) over comparable genes (Figure S1Q). The slight reduction of detected genes at the lowest input (∼10^4^ cells) may originate from the shallow complexity of the transcripts in the lysate. In addition to the ORF-wise analysis, codon-wise examinations of averaged ribosome occupancy showed high consistency for low-input data (Figure S1R). The comparable footprint distribution along the ORF was exemplified for *EEF2* mRNA (Figure S1S). These data indicated that the steps after RNase digestion do not hamper Thor-Ribo-Seq translatome analysis for small inputs.

Then, we conducted the whole Thor-Ribo-Seq procedure, including RNase digestion and downstream library preparation, from the low material inputs (denoted as “aliquoted-then-digested”) and assessed the bias derived from all steps (Figure S1I). As observed in the “digested-then-aliquoted” condition, the libraries in low-input samples were successfully prepared (Figure S1J-K) and showed the conventional hallmarks of Ribo-Seq (Figure S1T-W) and high correspondence in ORF-wise (Figure 1D) and codon-wise (Figure 1E-F) assessments to the data obtained from a standard amount of materials. We noted that the suppression of read duplications generated by T7 polymerase-mediated RNA amplification benefited the adequate evaluation of data. Our Thor-Ribo-Seq library contained UMIs at two distinct positions: at the 3′ end of footprints introduced by the first linker and at the 5′ end of footprints originating from the second linker (Figure S1B and S1X). Given the timing of the UMI addition, the 3′-end UMI suppressed the read duplications that occurred in the *in vitro* transcription, and the combination of the 3′ and 5′ UMIs limited read duplications during PCR (Figure S1X). We tested the potency of the UMIs to restrain the biases and found that the minimization of overamplified reads by *in vitro* transcription led to a high correlation among Thor-Ribo-Seq data compared to the same correction of biases in PCR (Figure S1P and S1Y). Our analyses mentioned above used 3′ UMI suppression unless noted.

We concluded that Thor-Ribo-Seq is a useful setup for translation surveys, tailored for low materials.

### High-throughput setup of Thor-Ribo-Seq in a semi-automated manner

Harnessing the versatility to handle small inputs, we developed a high-throughput platform based on a 96-well plate format. Given that ribosome-footprint complex isolation through sucrose cushion ultracentrifugation is a barrier for sample number scalability (Figure 2A), we instead implemented size-exclusion gel filtration resin (Sephacryl S400) on a 96-well plate with the filter at the bottom (Figure 2B), as conducted in the spin column of the resin ^8,39,40^. Here, large complexes such as the ribosome-footprint complex passed through the resin on the top filter plate and were collected on the bottom plate. We tested the impact of this ribosome purification and the downstream sample handling (the combination of TRIzol-mediated RNA purification and Ribo-FilterOut ^41^) on the quality of the data in both medium (∼10^5^ cells) and low (1.65 × 10^4^ cells, corresponding to ∼40% confluency in a 96-well culture) sample load (Figure S2A-M). Considering that the number of usable reads (*i.e.*, genome-mapped reads unmatched to non-coding RNAs such as rRNAs after UMI deduplication) and periodicity score ^42^ — a value indicating codon resolution, we ultimately choose the combination of gel-filtration and TRIzol purification since this condition provides a good balance of those two metrics (Figure S2D and S2G).

**Figure 2.**
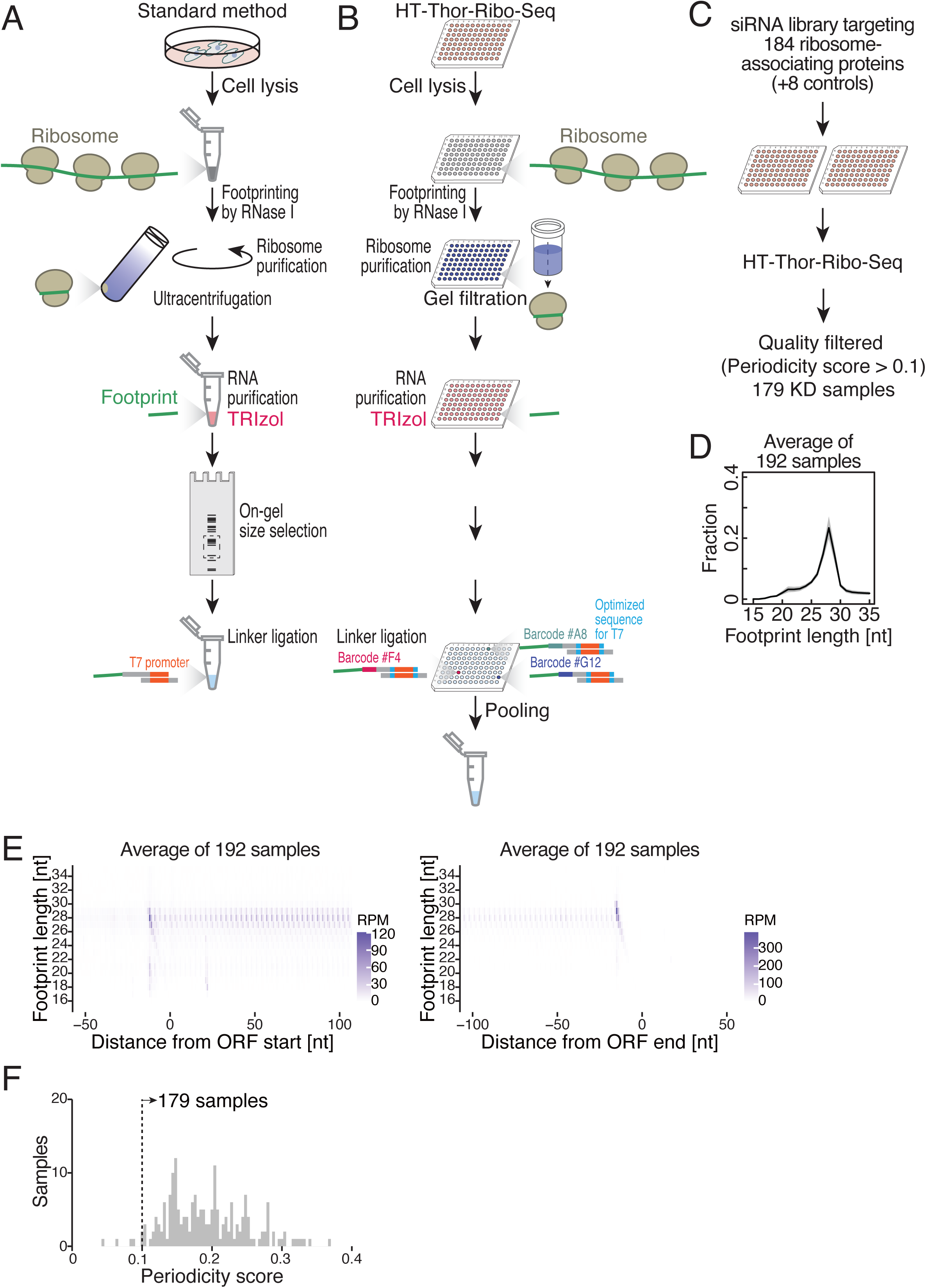
HT-Thor-Ribo-Seq captures the translation status from hundreds of samples in a parallel manner. (A and B) Schematic representation of the library preparation strategy of standard Ribo-Seq (A) and HT-Thor-Ribo-Seq (B). (C) Schematic representation of HT-Thor-Ribo-Seq for siRNA knockdown samples. (D) The distribution of ribosome footprint length for HT-Thor-Ribo-Seq of siRNA-treated samples. The average across 192 samples is shown as a black line. The shaded area represents ±1 standard deviation (SD). (E) Metagene plots of the 5′ end position of ribosome footprints around start (left) and stop (right) codons. The average across 192 samples was shown. X axis, the position relative to the start codon (0 as the first nucleotide of the start or stop codon); Y axis, footprint length; color scale, read abundance. RPM, reads per million mapped reads. (F) Histogram of periodicity scores for 192 samples. Samples with periodicity score > 0.1 (dashed line) were used for downstream analyses. KD, knockdown. See also Figures S2 and S3 and Tables S1, S5, and S6.

We redesigned linker oligonucleotides to implement the optimized DNA sequence upstream and downstream of the T7 promoter to maximize transcription ^43^ and sample barcodes for multiplexing 96 samples.

### Landscape of the translation regulatory network by ribosome-associated proteins

To benchmark the performance of high-throughput (HT)-Thor-Ribo-Seq, we applied it to the parallel knockdown of genes potentially related to translation regulation. Given the previous quantitative proteomics of HeLa and HEK293 cells with sucrose density gradient ultracentrifugation and extensive fractionations ^44^, we chose 184 ribosome-associated proteins (Table S1); this list included genes for which the function in translation has been reported, as well as genes whose function in translation regulation remains elusive. To understand the function of these ribosome-associated proteins in translation, we individually knocked them down by siRNAs in HEK293 cell culture on 96-well plates (Figure 2C). Including 8 samples with control siRNA treatment, HT-Thor-Ribo-Seq was performed for 192 conditions in total (*i.e.*, two 96-well plates). Simultaneously, we performed RNA-Seq to measure translation efficiency (footprint counts normalized by RNA-Seq counts). Notably, RNA-Seq confirmed the knockdown of target genes (Figure S3A).

Ribosome footprints from the HT-Thor-Ribo-Seq library were successfully obtained from all the wells on the two plates (Figure S3B-E) and showed hallmarks of ribosome footprints: two peaks at 22 and 28 nt in read distribution (Figure 2D) and triplet periodicity on the ORF (Figures 2E and S3F). The read counts on ORFs in control siRNA samples were highly correlated (Figure S3G), confirming the high reproducibility of data in this format. For quality check, we screened the sample based on the periodicity score (Figure 2F) and ultimately used 179 samples with high scores (> 0.1) for subsequent analysis (Table S1). We note that the low periodicity score may be associated with low read depth (Figure S3H).

The large-scale Ribo-Seq data allows a systematic comparison of the knockdown effects and functional estimation of ill-characterized proteins in translation. The 179 genes analyzed in this dataset span a wide range of functions, localizations, and domain structures (Figure 3A and Table S2, based on the Uniprot database). Here, considering changes in footprint, RNA, and translation efficiency, we performed clustering of the genes whose knockdowns have similar effects (Figures 3B and S3I-J). Indeed, this analysis aligned well with the knowledge of translational controls mediated by RBPs. In translation efficiency changes, the knockdown of two paralogs of cytoplasmic poly(A)-binding proteins (PABPs) (PABPC1 and PABPC4) and PABP-interacting PURA ^45^ and ATXN2 ^46^ showed similar impacts (Figure 3B). The same trends were held true in footprint changes (Figure S3I). Similarly, the depletion of proteins in the RNA silencing pathway, including AGO2 — a core protein of RNA-induced silencing complex (RISC) ^47^, TARBP2 (also known as TRBP) — a microRNA (miRNA) biogenesis factor ^48^, and STAU2 — a subcellular localization regulator of RISC ^49^, led to similar perturbations in translation efficiencies (Figure 3B).

**Figure 3.**
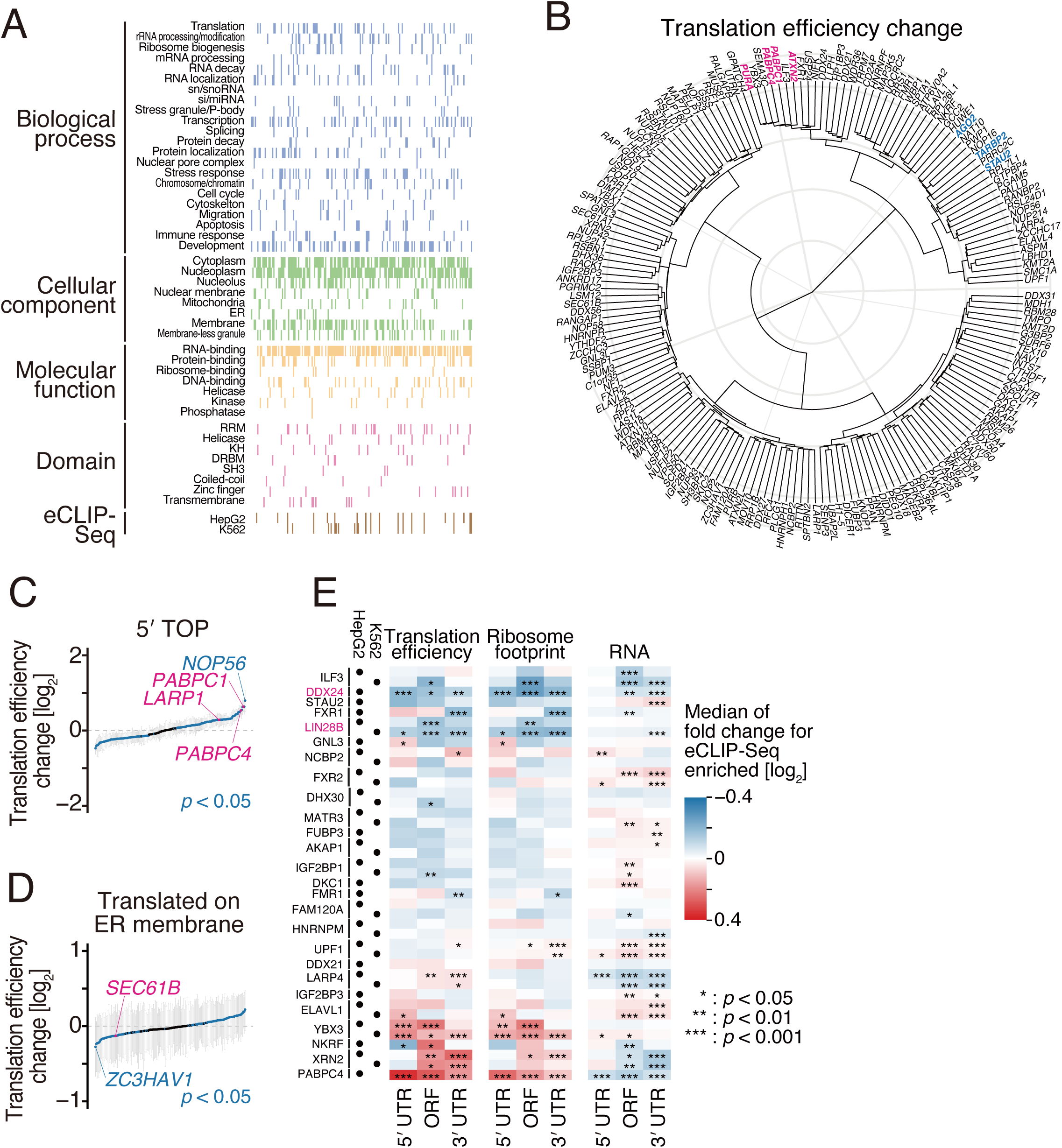
HT-Thor-Ribo-Seq reveals the global landscape of translation change for 179 siRNA knockdown samples. (A) Annotation of 179 knocked-down genes. Gene ontology terms (biological process, cellular component, and molecular function) and protein domains were assigned based on the UniProt database. The availability of eCLIP-Seq data deposited in the ENCODE project is also indicated. (B) Dendrogram of translation efficiency changes in 179 siRNA knockdown samples that passed the quality check threshold, relative to control samples. Knocked-down genes mentioned in the main text are highlighted. (C and D) Translation efficiency changes of genes with a 5′ TOP motif (C) and translated on the ER membrane (D) in 179 siRNA knockdown samples. The median (dot) and first and third quartiles (ticks) are shown. Knockdown samples showing significant up- or down-regulation of indicated mRNAs (*p* < 0.05, two-tailed Mann–Whitney *U* test with Benjamini-Hochberg adjustment) and samples mentioned in the main text are highlighted in blue and magenta, respectively. (E) Heatmap of translation efficiency, ribosome footprint, and RNA changes for mRNAs identified as eCLIP-Seq-enriched following siRNA knockdown of the corresponding gene. Enriched mRNAs were defined as those with log_2_(eCLIP-Seq enrichment) greater than 2 on the 5′ UTR, ORF, or 3′ UTR, as determined from ENCODE eCLIP-Seq data. Samples mentioned in the main text are highlighted in magenta. The color scale indicates the degree of median fold change in read counts. *, *p* < 0.05; **, *p* < 0.01; ***, *p* < 0.001, two-tailed Mann–Whitney *U* test with Benjamini-Hochberg adjustment. See also Figure S3 and Tables S1, S2, and S5.

Given that the sequence motifs on mRNAs play essential roles in the selective translation regulation, we next focused on mRNAs carrying well-known regulatory motifs such as 5′ tandem oligopyrimidine (TOP) sequence just after the m^7^G cap ^50^ (Figure 3C). Our translation efficiency measurement recapitulated the previous reports that described inhibitory mechanisms of translation: LARP1, a direct binder to m^7^G cap and the TOP motif, represses the translation of TOP mRNAs together with PABPC1 ^51^ (Figure 3C). These protein depletions induced the translational depression of TOP mRNAs (Figure 3C). Noteworthy, we found other proteins that showed more prominent impacts than the known factors (*e.g.*, NOP56), suggesting that they may play important roles in the translational regulation through this motif.

As translation efficiency is a metric to normalize RNA abundance changes, the effect on RNA level was not included. We note that a similar analysis on RNA changes directly again showed good alignment with the known regulations through RBPs and RNA elements. This included examples that LARP1 and PABPs stabilize TOP mRNAs ^52–54^ (Figure S3K).

Translation often occurs locally, such as on the endoplasmic reticulum (ER) membrane. Compared to the proximity ribosome profiling data for the ER membrane ^55^, we confirmed that the depletion of translocon SEC61B reduced translation efficiency of mRNAs translated on the ER (Figure 3D), whereas the depletion of this gene provided a minimal effect on RNA level (Figure S3L). We note that ZC3HAV1 (also known as ZAP) has a drastic role in ER translation. This may be consistent with the localization of ZC3HAV1 at the ER and the role in unfolded protein response (UPR) in ER stress ^56^.

Integration of published databases provided a deeper understanding of regulation mechanisms. Given the various RNA-binding proteins (RBPs) in the list (Figure 3A), we compared our data with the in-cell RBP-RNA binding spectrum investigated by enhanced crosslinking and immunoprecipitation followed by sequencing (eCLIP-Seq) in the Encyclopedia of DNA Elements (ENCODE) project ^57^. Throughout the knockdown Ribo-Seq data for 26 RBPs, we calculated the footprint, RNA, and translation efficiency changes of mRNAs associated with corresponding RBPs, employing the eCLIP-Seq dataset (Figure 3E). The positive correlation suggested that the RBP functions may play a role as an inhibitory factor, whereas the negative correlation was inferred as a positive factor. This analysis confirmed LIN28B as an activator (Figures 3E and S3M). Additionally, we found DDX24 as a potential translation activator (Figures 3E and S3M).

#### Ribosome traversal on 3′ UTR induced by the depletion of ribosome-associated proteins

Considering ribosome footprints outside of ORFs, we found a factor that affects ribosome traversal along mRNA after the completion of protein synthesis. Our quality check of the read distribution, which is typically enriched in ORFs, identified that *GNL3* knockdown led to read enrichment on the 3′ untranslated region (UTR) (Figure 4A). Metagene analysis around the stop codon confirmed this tendency (Figure 4B). Given that the 3′ UTR footprints did not show 3-nt periodicity (Figure 4C), the results indicated the accumulation of non-actively elongating ribosomes or unrecycled ribosomes after peptidyl-tRNA cleavage by termination factor eRF1, as reported in recycling factor ABCE1 (Rli1 in yeast) depletion ^58–61^. GNL3 has been characterized as a nucleolar pre-60S biogenesis factor with GTPase activity ^62,63^. However, the knockdown of ribosome biogenesis factors did not always show a similar effect in translation efficiency change (also footprint and RNA changes) as GNL3 knockdown (Figure S4A). Moreover, the knockdown of homologous protein GNL3L and other energy-dependent nucleolar pre-60S biogenesis factors (ATPase DDX18 and GTPase GTPBP4) did not echo the same effects (Figures 4B and S4B). These data indicated that the defect in ribosome recycling by GNL3 depletion was unlikely due to a global ribosome biogenesis defect, suggesting that recycling-deficient “scarred” ribosomes may be generated by GNL3 deficiency specifically.

**Figure 4.**
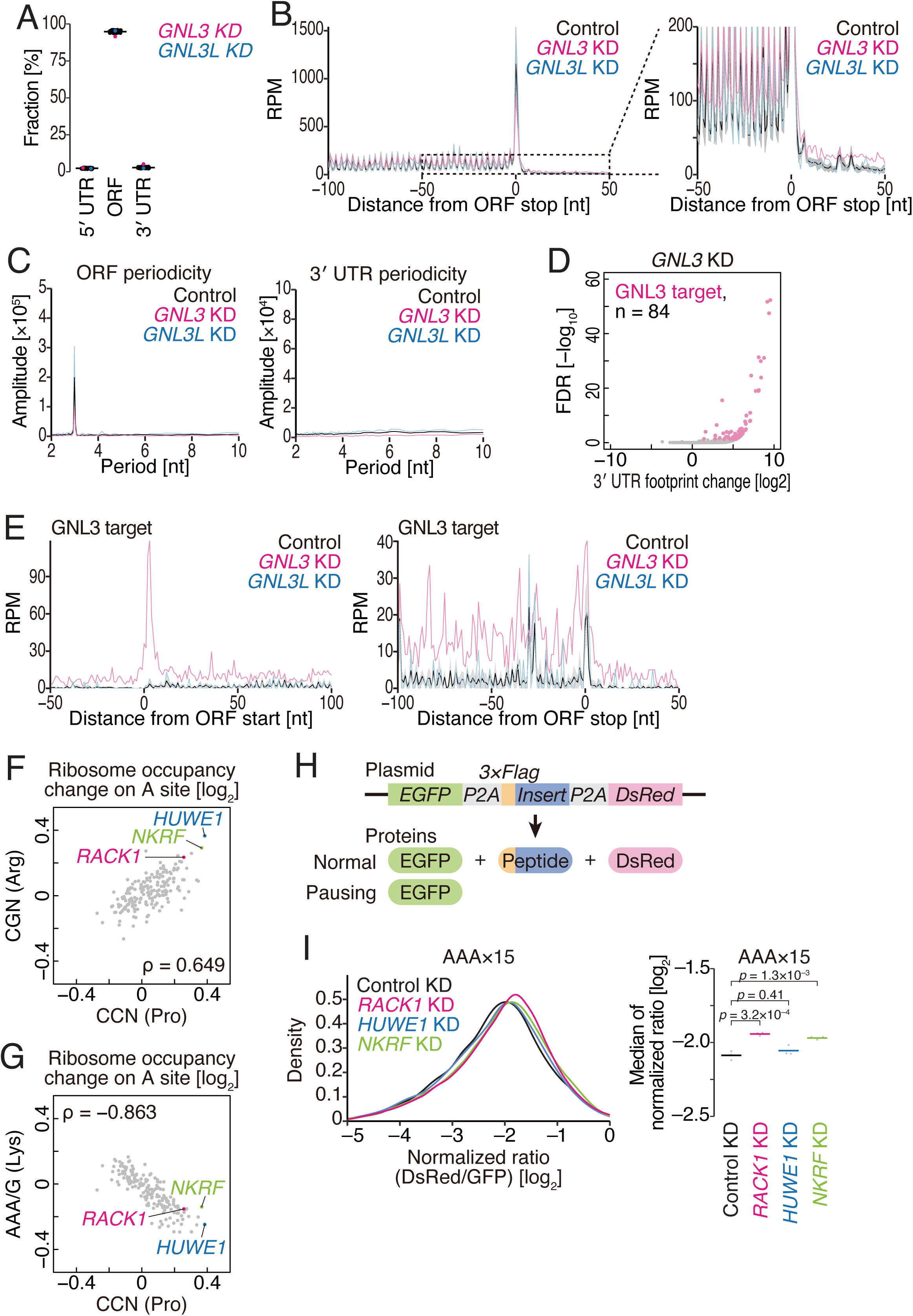
Identification of modulators of ribosome traversal. (A) Fraction of ribosome footprints on the 5′ UTR, ORF, and 3′ UTR. The indicated gene knockdown samples are color-highlighted. (B) Metagene plots of the 5′ end position of ribosome footprints around stop codons in the indicated knockdown samples. An expanded view is shown on the right. The indicated gene knockdown samples are color-highlighted. For control knockdown samples, the average across 8 samples is shown as a black line, and the shaded area represents ±1 standard deviation (SD). (C) Discrete Fourier transform of ribosome footprints mapped to the last 200 nt of the ORF region (left) and the first 200 nt of the 3′ UTR region (right). The indicated gene knockdown samples are color-highlighted. (D) Volcano plot of 3′ UTR footprint changes in *GNL3* knockdown. mRNAs possessing high 3′ UTR footprints [a false discovery rate (FDR) < 0.05] upon *GNL3* knockdown are highlighted and defined as “GNL3 target” mRNAs. The number (n) of GNL target mRNAs is shown. (E) Metagene plots for GNL3 target mRNAs (defined in D) showing the 5′ end positions of ribosome footprints around start (left) and stop (right) codons. For control knockdown samples, the average across 8 samples is shown as a black line, and the shaded area represents ±1 standard deviation (SD). (F and G) Scatter plot of changes in ribosome occupancy on CCN versus CGN codons (F) and CCN versus AAA/AAG codons (G) at the A site. Knocked-down genes mentioned in the main text are highlighted. ρ, Spearman’s rank correlation coefficient. (H) Schematic representation of the reporter assay for ribosome stalling. (I) Density plot (left) and median values (right) of the normalized ratio between DsRed and GFP signals for the indicated knockdowns. The *p* values were calculated by the two-sided Tukey-Kramer multiple comparison test. KD, knockdown; RPM, reads per million mapped reads. See also Figures S4 and S5 and Table S5.

The GNL3-mediated recycling deficiency was mRNA-selective (Figure 4D). Among the 84 genes classified as 3′ UTR read-upregulated genes in *GNL3* knockdown (Figure 4D, GNL3 target), we observed strong accumulation of ribosome footprints not only on the 3′ UTR but also on the 5′ UTR and ORF regions by GNL3 depletion (Figure 4E). For these genes, ribosome footprints on the ORF were longer and less periodic in *GNL3* knockdown cells (Figures S4C-D). In contrast, the other genes showed regular ribosome footprint distribution along mRNAs (Figure S4E) and conventional read length (Figure S4F) even with *GNL3* knockdown. As footprints generally originate from 80S, our results suggest that ribosomes generated under GNL3 loss are defective not only in recycling but also in precise initiation processes, leading to aberrant translation in an mRNA-selective manner.

### Ribosome stalling by the depletion of ribosome-associated proteins

In addition to ORF-wise analysis, codon-wise ribosome occupancy was further assessed in HT-Thor-Ribo-Seq data. Control samples showed a high correspondence of ribosome occupancy on either A-, P-, or E-sites (Figure S5A-C), confirming the reproducibility in the high-throughput format. Clustering analysis revealed that a subgroup of samples increased ribosome occupancy on all 4 Pro codons (CCN) and 4 out of 6 Arg codons (CGN) at the A site (Figure S5D, cluster 6), but reduced it on Lys codons (AAA/AAG). These effects were not observed at the P and E sites (Figure S5E-F). The effects on CCN Pro codons were correlated with CGN Arg codons and anti-correlated with Lys codons in the knockdown samples (Figure 4F and 4G), suggesting the same mechanism behind the ribosome stalling on both types of codons. Ribosome stalling on CSN (where S represents C and G) codons was remarkable in RACK1 (a component of 40S required for the ribosome stalling resolution) ^64–66^, HUWE1 (an E3 ubiquitin ligase for orphan ribosome proteins) ^67^, and NKRF (a ribosome biogenesis factor) ^68,69^ (Figures 4F-G and S5D). Notably, these knockdowns were not associated with the downregulation of eIF5A, which facilitates translation elongation on the Pro-Pro motif (Figure S5G) ^70^.

Given that poly-lysine (especially AAA codon repeat) has been a well-described inducer of ribosome-associated quality control (RQC) and that RACK1 contributes to the RQC ^66,71–74^, forming a unique interface in collided ribosomes ^75–77^ required for RQC induction, we tested whether the factors found in our screen play a role in RQC. For this purpose, we conducted flow cytometry-based reporter assays ^66,78^ by inserting test sequences between GFP and DsRed and liberating the GFP, poly-lysine peptide, and DsRed as individual proteins by viral P2A sequences ^79^ (Figure 4H). Aligning with earlier studies ^77,78,80,81^, AAA repeat insertion hampers ribosome elongation (Figure S5I). As previously reported ^66,74^, RACK1 depletion induced the readthrough of the AAA codon stretch, as represented by a higher DsRed/GFP ratio (Figure 4I). As similar effects were observed in NKRF knockdown (Figure 4I), this factor may have a function to facilitate RQC.

### Large-scale assessment of ribosome collision by HT-Thor-Disome-Seq

The differential ribosome stalling by knockdown of ribosome-associated proteins led us to investigate the ribosome collision, where stalled ribosomes create a bottleneck for those following behind. Given that two queued ribosomes (or di-ribosomes/disomes) generate longer ribosome footprints, Disome-Seq surveys ribosome collision sites in a genome-wide manner and at high resolution ^30,82–85^. Here, we applied the Disome-Seq to our high-throughput format (HT-Thor-Disome-Seq). The data exhibited the typical characteristics of disome footprints: 52- and 59-nt peaks (Figure 5A) and strong accumulation at stop codons (Figure 5B), as reported previously ^30,84,85^. The aggregation of all samples recapitulated the known ribosome pause sites on *XBP1u* and *SEC61B* mRNAs ^30,86^ (Figure S6A). A high correlation of ORF-wise disome footprints among replicates ensured the reproducibility (Figure S6B).

**Figure 5.**
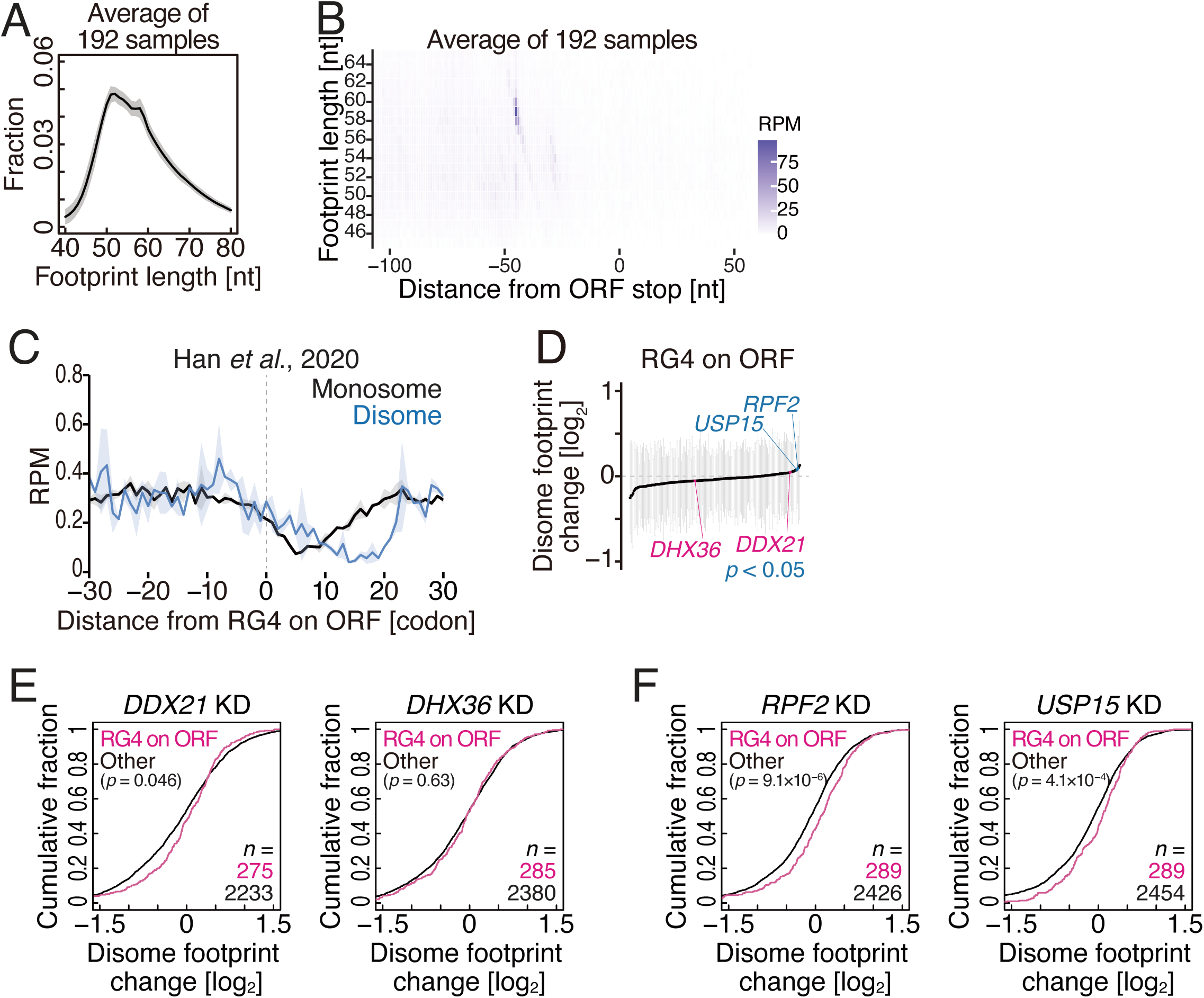
HT-Thor-Disome-Seq detects ribosome collisions for hundreds of samples in a parallel manner. (A) The distribution of footprint lengths for HT-Thor-Disome-Seq of siRNA-treated samples. The average across 192 samples is shown as a black line. The shaded area represents ±1 standard deviation (SD). (B) Metagene plots of the 5′ end position of disome footprints around stop codons in the indicated experiments. The average of 192 samples was shown. X axis, the position relative to the stop codon (0 as the first nucleotide of the stop codon); Y axis, footprint length; color scale, read abundance. (C) Metagene plot for disome footprint from the start site of RG4 on ORFs. The previous monosome and disome profiling data from ^30^ were analyzed. The averages across duplicates are shown as thick lines, and the shaded areas represent ±1 standard deviation (SD). (D) Disome footprint changes of genes with RG4 on ORFs. The median (dot) and first and third quartiles (ticks) are shown. Knockdown samples showing significant up- or down-regulation of indicated mRNAs (*p* < 0.05, two-tailed Mann–Whitney *U* test with Benjamini-Hochberg adjustment) and samples mentioned in the main text are highlighted in blue and magenta, respectively. (E and F) Cumulative distributions of disome footprint changes in the indicated knockdowns. The *p* values were calculated with the two-sided Mann‒Whitney *U* test. The gene number (n) of each group is shown. KD, knockdown; RPM, reads per million mapped reads. See also Figure S6 and Tables S5 and S6.

Strong RNA secondary structures are suggested to induce ribosome stalling. We particularly focused on RNA G-quadruplex (RG4) since its strong potential for ribosome roadblock ^87–89^. Through the reanalysis of our previous Disome-Seq data ^30^ and the comparison with the recent genome-wide survey of RG4 (RG4-Seq 2.0) ^90^, we found that disome footprints were accumulated 5-9 codons upstream from the RG4 structure (Figure 5C). This positional relationship between the paused ribosome and RG4 aligned well with the study ^87^. *LRRC59* and *ZNF593* mRNAs exemplified the association of ribosome collision and RG4 formation (Figure S6C). These data indicate that RG4 is a source of ribosome collision.

Given the harmful effects of RG4 for ribosome traversal, we reasoned that factors such as helicases may counteract RG4 formation for smooth ribosome traversal. Through HT-Thor-Disome-Seq data, we surveyed factors whose depletion increased disome footprints on RG4 containing ORFs (Figure 5D). Indeed, DDX21, a known RG4 helicase ^91–93^, was found as such a factor (Figure 5D and 5E). On the other hand, DHX36, another RG4 helicase ^94–97^, showed no obvious effect (Figure 5D and 5E). Possibly, the sequences in the loop and/or the flanking region of RG4 may determine the target RG4 specificity of helicases and steric hindrance ability for elongating ribosomes. Our screening suggested that RPF2 and USP15 enhance ribosome traversal over RG4 (Figure 5D and 5F).

### Survey for mitochondrial translation regulators

As standard Ribo-Seq ^11,41,98–102^, HT-Thor-Ribo-Seq also captured mitoribosome footprints and measures mitochondrial translation status (Figure 6A). To date, known regulators of mitochondrial translation have been limited to TACO1 ^103–105^, LRPPRC-SLIRP ^106–110^, C1QBP/p32 ^111^, MITRAC15 ^112^, PREPL ^113^, TRAP1 ^114^, DHX30 ^115^, TMEM223 ^116^, and IGF2BP1 ^115^. All of these factors function as positive regulators. However, to our knowledge, no mitochondrial translation repressors have yet been identified in humans.

**Figure 6.**
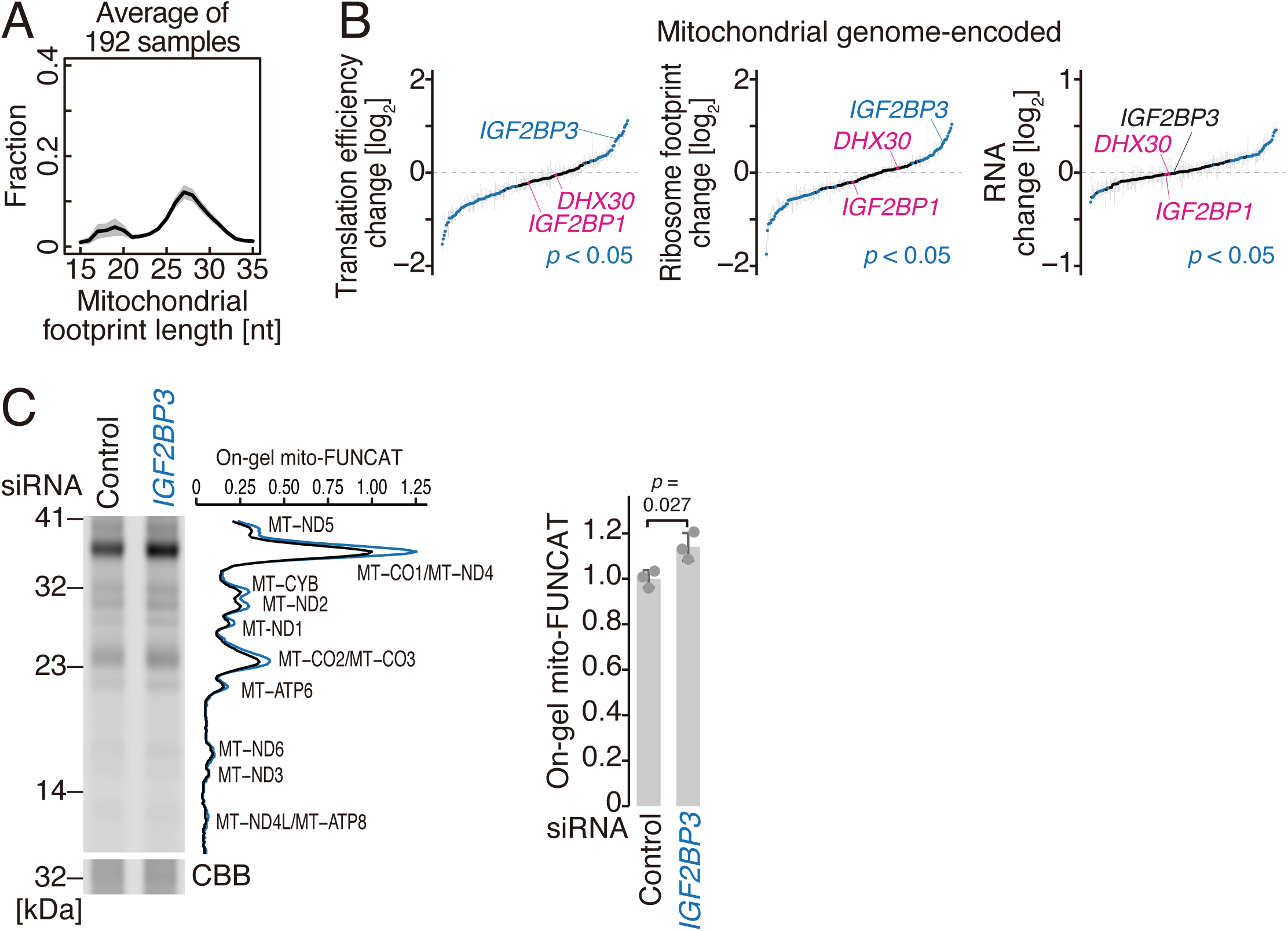
HT-Thor-Ribo-seq surveys mitochondrial translation regulators. (A) The distribution of mitochondrial footprint length for HT-Thor-Ribo-Seq of siRNA-treated samples. The average across 192 samples is shown as a black line. The shaded area represents ±1 standard deviation (SD). (B) Translation efficiency (left), ribosome footprint (middle), RNA (right) changes of mitochondrial genome-encoded genes in 179 siRNA knockdown samples. The median (dot) and first and third quartiles (ticks) are shown. Knockdown samples showing significant up- or down-regulation of indicated mRNAs (*p* < 0.05, two-tailed Mann– Whitney *U* test with Benjamini-Hochberg adjustment) and samples mentioned in the main text are highlighted in blue and magenta, respectively. (C) On-gel FUNCAT experiments of mitochondrial translation after *IGF2BP3* knockdown. Quantification of band intensities along the lanes (middle) and total intensities [right, normalized by the total protein load measured by Coomassie Brilliant Blue (CBB) staining] is shown. On the right panel, the means (bars), SDs (errors), and individual replicates (points, n = 3) are shown. The *p* value was calculated by the two-sided Student’s t-test. See also Tables S5 and S6.

Direct measurement of mitochondrial translation changes for all 13 ORFs encoded in the mitochondrial genome ^117^ in siRNA screening revealed the negative translation regulators, in addition to the translation activators. Our data confirmed that IGF2BP1 promotes mitochondrial translation ^115^, as its knockdown reduced the mitochondrial translation efficiency (Figure 6B). We found 39 potential negative regulators of mitochondrial translation (*p* < 0.05 by Mann-Whitney *U* test with Benjamini-Hochberg adjustment), as we found increased mitochondrial protein synthesis by the knockdown of the factors. Interestingly, we observed that the knockdown of the IGF2BP3, a paralog of IGF2BP1, conversely showed upregulation of mitochondrial translation. On-gel mitochondrial-specific fluorescent noncanonical amino acid tagging (on-gel mito-FUNCAT) ^118,119^ further validated this result (Figure 6C). Although this protein was shown to associate with mitoribosomes ^115^, the molecular role has remained elusive. Our study indicated IGF2BP3 as a repressor of mitochondrial translation.

### HT-Thor-Ribo-Seq reveals the symptom severity-dependent alteration of translation in COVID-19 patients

This high-throughput approach to translation led us to investigate patient-derived samples, which may promote the understanding, diagnosis, and therapeutic strategies of human diseases. Here, we applied HT-Thor-Ribo-Seq to PBMCs, in which T cells occupy the major cell population, from severe acute respiratory syndrome coronavirus 2 (SARS-CoV-2)-infected patients (collected from March 2020 to December 2021 in Japan). We isolated PBMC from 93 patients with various severities and three non-infected people (Figure 7A and Table S3). Ribosome footprints from PBMCs again showed typical distribution in length (Figure S7A) and 3-nt periodicity along the ORFs (Figure S7B). We also performed the RNA-Seq from the same lysates. In both types of data, high reproducibility of the data was observed via healthy persons’ samples (*n* = 3) (Figure S7C). Through the curation based on periodicity scores and gene numbers detected in Ribo-Seq (Figure S7D-G), we ultimately focused on 70 patients’ and 3 non-infected persons’ samples for downstream analysis.

**Figure 7.**
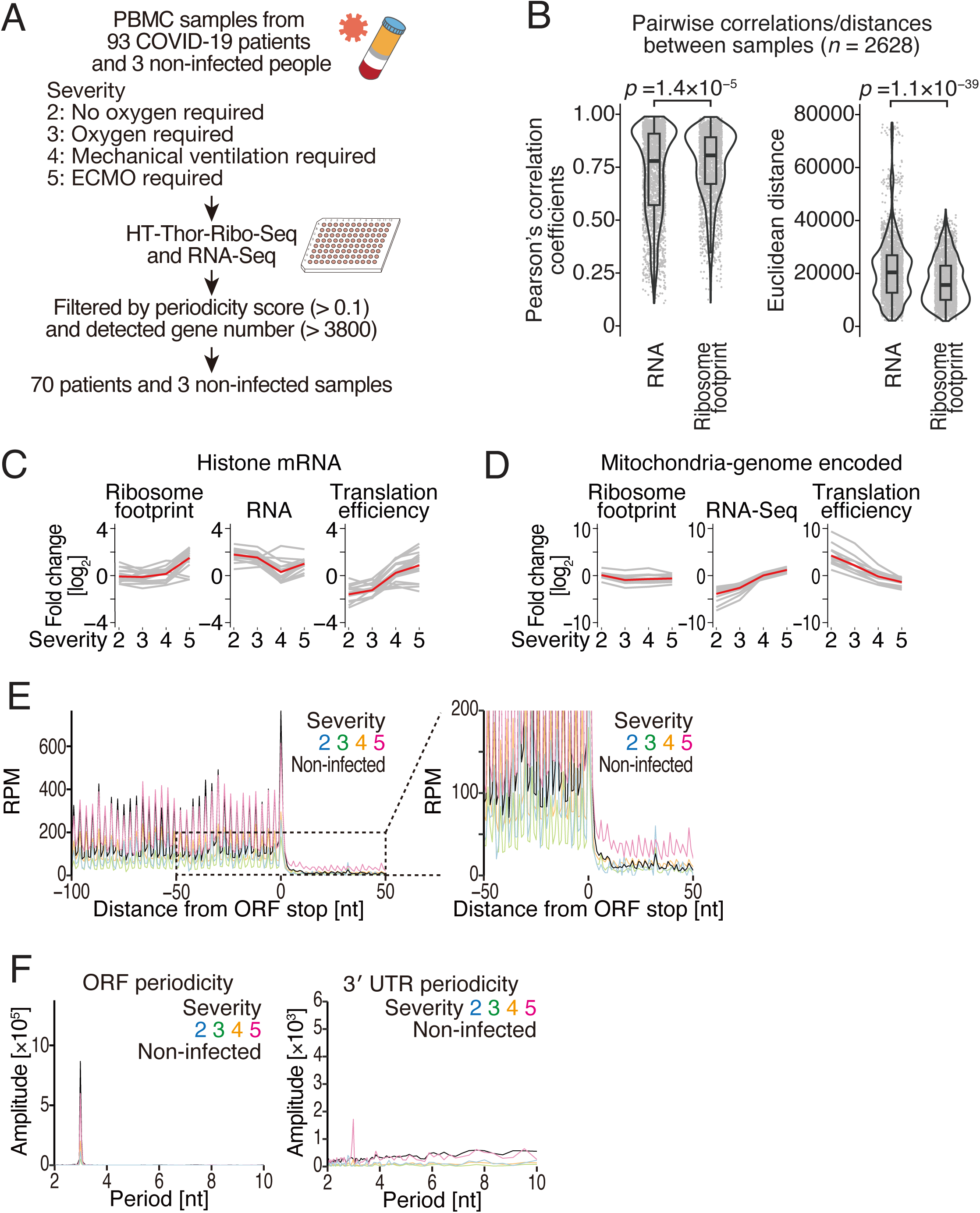
HT-Thor-Ribo-Seq reveals translation landscape of COVID-19 patients. (A) Schematic representation of HT-Thor-Ribo-Seq for PBMC samples from COVID-19 patients. (B) Violin plots of pairwise Pearson’s correlation coefficients (left) and Euclidean distances (right) between samples. Read counts were TPM (transcripts per kilobase million)-normalized. Box plots show the median (centerline), upper/lower quartiles (box limits), and 1.5× interquartile range (whiskers). The *p* values were calculated with the two-tailed Mann‒Whitney *U* test. The number (n) of all combinations between 73 samples (_73_C_2_) is shown. (C and D) Ribosome footprint, RNA, and translation efficiency changes of histone genes (C) and mitochondrial genome-encoded genes (D) for each severity group. The mean fold change for each group is depicted as a red line. (E) Metagene plots of the 5′ end position of ribosome footprints around stop codons in the indicated samples. An expanded view is shown on the right. (F) Discrete Fourier transform of ribosome footprints mapped to the last 200 nt of the ORF region (left) and the first 200 nt of the 3′ UTR region (right). RPM, reads per million mapped reads. See also Figure S7 and Tables S3, S4, S5 and S6.

To globally survey the impact of transcriptome and translatome among the patient samples, here we calculated Pearson’s correlation coefficients and Euclidean distance of all the data pairs in each dataset. In both metrics, we found that Ribo-Seq sets generally showed less variability than RNA-Seq data sets (Figure 7B), suggesting the buffering of transcriptional fluctuation at the translational level.

To ensure SARS-CoV-2 infection in the data, we assessed genes whose expression changes in COVID-19 infection were reported. This analysis included ferritin (*FTL* and *FTH1*), thrombospondin (*THBS1*), and transcription factor *MAFB* ^120–123^. We observed high RNA levels of those genes in RNA-Seq and the reflected footprints in Ribo-Seq (Figure S7H). In addition, especially in *FTL* and *FTH1*, translation efficiency was also increased, indicating translational impacts during viral infection (Figure S7H).

Beyond the previously described genes, we observed widespread translation efficiency changes across the transcriptome in patients (Figure S7I). To identify genes with modulated translation efficiency along with symptom severity, we classified samples by artificial respiration methods at the time of sampling. Clustering analysis (Figure S7J-K) defined genes with increased translation efficiency (cluster 2) in a severity-dependent manner. The cluster 2 genes were characterized by histone genes (Table S4), as highlighted by the enriched gene ontology (GO) term in heterochromatin organization (Figure S7L). Indeed, most histone genes showed drastic increases in translation efficiencies with high severity (Figure 7C).

Conversely, a subset of mRNAs in cluster 8 showed reduced translation efficiency (Figure S7J-K). These genes were associated with mitochondrial functions, as presented by related GO term enrichment (Figure S7L). Therefore, we reasoned that mitochondrial translation may be affected in SARS-CoV-2 patients, as our HT-Thor-Ribo-Seq detects mitochondrial footprints (Figure 6). Indeed, translation efficiencies of all 13 ORFs encoded in the mitochondrial genome were decreased along with symptom severity (Figure 7D). Instead, possibly due to compensation mechanisms, mitochondrial mRNA abundance was increased (Figure 7D). Given that all 13 ORFs in the mitochondrial genome encode subunits of the OXPHOS complex, we wonder about the influence on the subunit supply from the cytosol. However, translation from mRNAs for nuclear genome-encoded OXPHOS complex subunits (Figure S7M), or more globally, mitochondria-targeted proteins (Figure S7N), was not affected by symptom severity. Thus, our data revealed the specificity of symptom-associated defects of mitochondrial translation within the matrix. This molecular phenotype could be explained by an increased population of exhausted T cells ^124,125^, which are tightly associated with mitochondrial impairment ^126–128^. Alternatively, considering mitochondrial translation is promoted by fever during infection and inflammation ^129^, the reduced efficiency of mitochondrial translation may be detrimental for the viral response.

In addition to the ORF-wise alterations, codon-wise ribosome occupancies were modulated after infection (Figure S7O). Higher ribosome occupancies at the A-site on codons were found for Asp, Gly, and Val codons, implying the shortage of these amino acids, possibly due to the metabolic change in exhausted T cells ^130,131^.

We found the C-terminal extension of proteins associated with severe COVID-19 symptoms. Footprints emerged from 3′ UTR but not 5′ UTR were increased in a severity-dependent manner (Figures 7E and S7P). The 3′ UTR footprints showed 3-nt periodicity (Figure 7F), indicating stop codon readthrough. This was in stark contrast to the molecular phenotype found in *GNL3* knockdown in HEK293 cells (Figure 4C).

We also categorized genes with RNA abundance changes. Subgroups of mRNAs were increased in their abundance in high severity (RNA clusters 4 and 5) (Figure S7Q-R). These mRNAs encoded translation machinery in terms of GO (Figure S7S). Since mRNAs encoding ribosome proteins and translation regulators possess the TOP motif ^50^, we focused on this mRNA group. Indeed, the TOP mRNAs showed an increased RNA, reflecting the translation level observed in Ribo-Seq (Figure S7T). This may be due to the mRNA stabilization through the LARP1 binding to this motif ^52,132^, which should be enhanced in mTORC1 inhibition. Consistent with this, attenuation of mTORC1 activity has been known in exhausted T cells by chronic virus infection ^133^. We note the limited detection of reduced translational efficiency in TOP mRNAs (Figure S7T).

In conclusion, these observations provide a hallmark of ribosome traversal dysregulation associated with severe symptoms of SARS-CoV-2. The application of HT-Thor-Ribo-Seq to patient samples enables comprehensive monitoring of translational fluctuation induced by diseases.

## Conclusion

This study developed Thor-Ribo-Seq as a sensitive methodology to perform translatome analysis with limited materials and a high-throughput derivative for translational surveys on massive parallel samples. RNA amplification with T7 RNA polymerase-mediated *in vitro* transcription, which lies at the core of this improvement, has been a useful option for handling low amounts of nucleic acids ^134,135^. Because of the linear amplification, this strategy has been reported to be a less biased approach ^136^ than exponential expansion by PCR. In practice, linear amplification has been implemented in diverse single-cell sequencing techniques, such as whole-genome analysis ^137^, RNA-Seq ^138,139^, and epigenomic profiling ^140,141^. Similarly, the investigation of limited transcriptomes on RNA-binding proteins was greatly improved by this approach ^142,143^. However, all those methods required the formation of double-stranded DNA (dsDNA) for *in vitro* transcription; thus, the execution of this technique was possible only at later steps of library preparation. In contrast, our method using single-stranded RNAs as templates has a strong benefit since its execution at an early step of the procedures effectively counteracts material loss. Generally, this technique should be easy to implement in library construction in other types of RNA-based sequencing technologies, including translation complex profile sequencing (TCP-Seq or 40S footprinting) ^144–148^, CLIP-Seq ^149^, and tRNA-Seq ^150^, all of which handle limited amounts of RNA fragments.

In addition to library construction, our library preparation strategy also has an advantage in computational analysis. Our strategy should outperform A-site offset assignments compared to preexisting methods for small inputs. Micrococcal nuclease (MNase), which has been used for scRibo-Seq ^10^ due to the stringent activity control by Ca^2+^ ions, has an A/U preference ^151^. The poly(A) tailing and template switching used in the one-pot reaction ^9,11–13^ add homopolymeric nucleotides to the footprints and thus generate ill-defined read boundaries at both the 5′ and 3′ ends. These drawbacks hamper the A-site offset estimation in ribosome footprints. In contrast, our strategy uses less biased RNase I and double-linker ligation to explicitly determine the ends of reads, preventing ambiguity regarding the A-site position reference.

Ultimately, HT-Thor-Ribo-Seq provides a widely applicable “screening” platform to monitor translational impacts, potentially by small molecule compounds, gene knockdown/knockout/overexpression, pathogenic variants, and beyond. Even via this work, the translational surveys revealed the unexpected dynamics of protein synthesis and simultaneously raised new questions (how the ribosome-associated factors control the initiation, elongation, and ribosome collision, how the viral infection causes the dysregulation, etc). Given that a collection of large Ribo-Seq datasets generates a machine learning model to predict translation efficiency and identifies a translational covariation associated with cell types ^152,153^, HT-Thor-Ribo-Seq should further boost these computational approaches. Our study paves the way for the future expansion of translational regulations in diverse contexts.

## Acknowledgments

We are grateful to all the members of the Iwasaki laboratory (RIKEN), Mara Anais Llamas-Covarrubias (National Institute of Biomedical Innovation), Ryuta Uraki (The University of Tokyo), Yasushi Kogo (RIKEN), and Taiki Baba (Nagasaki University), for constructive discussions, technical help, and critical reading of the manuscript. The COVID-19 clinical study was supported by Japan’s SIP “AI Hospital” program in FY2021. Clinical specimens were collected with the cooperation of Yuji Fujino (Osaka University Hospital), Nobuaki Shima (Hiroshima University Hospital), and Toru Kotani (Showa University Hospital). We also thank the HOKUSAI SailingShip supercomputer facility at RIKEN for the computational support and the Support Unit for Bio-Material Analysis, RIKEN CBS Research Resources Division, for Sanger sequencing and flow cytometry analysis. This work was supported by the Ministry of Education, Culture, Sports, Science and Technology (MEXT) (JP20H05784 and JP24H02307 to S.I.; JP21H05734, JP23H04268, and JP25H01440 to Y.S.), the Japan Agency for Medical Research and Development (AMED) (JP20gm1410001 to S.I.; JP23gm6910005 to Y.S.), the Japan Society for the Promotion of Science (JSPS) (JP23H02415 and JP23H00095 to S.I.; JP21K15023 and JP23K05648 to Y.S.; JP24K23198 to T.W.), the Nakajima Foundation (to Y.S.), the Exploratory Research Center on Life and Living Systems (ExCELLS) (23EXC601-2 and 25EXC602-4 to Y.S.), the National Institutes of Natural Science (NINS) (OML022505 to A.Y.), and RIKEN (Pioneering project “Biology of Intracellular Environments” to S.I. and Y.S.; RIKEN TRIP initiative “TRIP-AGIS” to S.I.; Incentive Research Project to T.W. and A.Y.).

## Author contributions

Conceptualization: Y.S., M.M., and S.I.;

Methodology: Y.S., M.M., Y.K., T.W., A.Y., Y.I., and S.I.;

Formal analysis: Y.S., M.M., Y.K., T.W., and A.Y.;

Investigation: Y.S., M.M., Y.K., T.W., and A.Y.;

Resources: Y.K., Y.M., and Y.I.;

Writing – Original Draft: Y.S. and S.I.;

Writing – Review & Editing: Y.S., M.M., Y.K., T.W., A.Y., Y.M., Y.I., and S.I.;

Visualization: Y.S. and T.W.;

Supervision: Y.S., Y.M., Y.I., and S.I.;

Project Administration: S.I.;

Funding Acquisition: Y.S., T.W., A.Y., and S.I.

## Experimental procedures

### Ethical statement

All procedures for a multi-center prospective cohort study on COVID-19 and PBMC collection were conducted with approval from the local ethics committees of the National Institute of Biomedical Innovation, Health and Nutrition (NIBIOHN), Osaka University Hospital, Hiroshima University Hospital, Showa University Hospital, and RIKEN.

### Cell lines

HEK293 Flp-In T-REx cells (Thermo Fisher Scientific, R78007) and HEK293 cells (ATCC, CRL-1573) were cultured in DMEM, high glucose, GlutaMAX Supplement (Thermo Fisher Scientific, 10566-016) or DMEM (4.5 g/l Glucose) with L-Gln and Sodium Pyruvate with 10% FBS at 37°C with 5% CO_2_, according to the manufacturer’s instructions. The cell culture was routinely tested for *Mycoplasma* contamination with the e-Myco VALiD Mycoplasma PCR Detection Kit (iNtRON Biotechnology, 25239) and confirmed to be negative.

### Peripheral blood samples from participants

We collected peripheral blood samples from healthy volunteers and COVID-19 patients who were admitted to general wards or intensive care units (ICUs) at Osaka University Hospital, Hiroshima University Hospital, and Showa University Hospital in Japan between March 1, 2020, and December 31, 2021, and tested positive for SARS-CoV-2 by PCR. This timeframe encompassed the first through fifth waves of the COVID-19 pandemic in Japan. Patient’s severity was classified in 4 levels as shown in Table S3 based on the criteria of the Ministry of Health, Labour and Welfare in Japan (Treatment guide for COVID-19 v5.2, https://www.mhlw.go.jp/content/000815065.pdf).

Peripheral blood samples were collected at multiple time points, including upon hospital admission and immediately before discharge, at intervals of approximately 1 to 2 weeks, depending on the patient’s clinical course. Among these, samples obtained during hospitalization were used in this study. In addition, samples from 3 healthy volunteers were included as controls. PBMCs were isolated using pluriMate II (pluriSelect Life Science, 44-10050-05) and stored at −80°C.

### Lysate preparation

#### HEK293 Flp-In T-REx cells and HEK293 cells

The cell lysate was prepared as described previously ^8^. Briefly, HEK293 Flp-In T-REx cells were cultured in a 10-cm dish, washed with ice-cold PBS, and then lysed with lysis buffer [20 mM Tris-HCl pH 7.5, 150 mM NaCl, 5 mM MgCl_2_, 1 mM dithiothreitol (DTT), 1% Triton X-100, 100 µg/ml cycloheximide, and 100 µg/ml chloramphenicol]. The extract was treated with 0.025 U/µl Turbo DNase (Thermo Fisher Scientific, AM2238), clarified by centrifugation at 20,000 × *g*, 4°C for 10 min, flash-frozen in liquid nitrogen, and stored at −80°C. The RNA concentration in the lysate was measured by a Qubit RNA BR Assay Kit (Thermo Fisher Scientific, Q10210). Of note, cells were treated with DMSO for 15 min in the standard Ribo-Seq and Thor-Ribo-Seq.

For siRNA knockdown experiments, 92 µl of 0.6 × 10^5^ cells/ml HEK293 cells were seeded onto each well of 96-well plates. The next day, the premixes of 0.5 µl of 5 µM siRNA (Cherry-pick library for ON-TARGETplus SMARTpool, Dhamacon, Table S1), 0.15 µl of *Trans*IT-X2 (Mirus Bio, MIR6003), and 9 µl of Opti-MEM (Thermo Fisher Scientific, 31985070) were added to each well (*i.e.*, 25 nM siRNA at the final concentration). After 72 h incubation, cells were washed with ice-cold PBS and lysed with 50 µl of lysis buffer with 0.025 U/µl Turbo DNase (Thermo Fisher Scientific, AM2238) by pipetting 3 times. Using an ASSIST PLUS pipetting robot (INTEGRA, 4505), the cell lysates were loaded on 0.2-µm filter plates (Corning, FiltrEX 3505) and layered on reservoir plates (Corning, AxyGen PCR-96-FS-C). After enclosing the top of the filter plates with sealers (Greiner, Bio-One 676051), layered plates were centrifuged at 1,500 × *g*, 4°C for 10 min by Sorvall Legend XFR (Thermo Fisher Scientific, 75004540) with HIGHPlate 6000 Microplate Rotor (Thermo Fisher Scientific, 75003606). The RNA concentrations of the lysates (∼40 µl captured in the reservoir plates) were measured by Qubit RNA HS Assay Kit (Thermo Fisher Scientific, Q32855). Typically, lysates contained ∼12 ng/µl of total RNA. After enclosing the top of the reservoir plates with PX1 PCR Plate Sealer (Bio-Rad, 1814000J1), the lysates on the plates were flash-frozen in liquid nitrogen and stored at −80°C.

#### PBMCs

Frozen stocks of PBMCs were thawed with thawing media (10% FBS and 100 µg/ml cycloheximide in RPMI-1640) at 37°C, washed with ice-cold PBS, and then lysed with lysis buffer. The extract was treated with 0.025 U/µl Turbo DNase (Thermo Fisher Scientific, AM2238), clarified by centrifugation at 20,000 × g, 4°C for 10 min, flash-frozen in liquid nitrogen, and stored at −80°C. The RNA concentration in the lysate was measured by a Qubit RNA BR Assay Kit.

### Library preparation for standard Ribo-Seq

Standard Ribo-Seq was conducted as described previously ^8^. Cell lysate containing 10 µg of RNA was brought to a volume of 300 µl by lysis buffer, treated with 20 U of RNase I (LGC Biosearch Technologies, N6901K) (*i.e.*, 0.067 U/µl) at 25°C for 45 min, placed on ice, supplemented with 200 U of SUPERase•In RNase Inhibitor (Thermo Fisher Scientific, AM2694), and ultracentrifuged in a sucrose cushion at 100,000 rpm at 4°C for 1 h with a TLA110 rotor (Beckman Coulter). The ribosome pellet was dissolved in TRIzol reagent (Thermo Fisher Scientific, 15596018), and RNA was isolated with Direct-zol RNA MicroPrep (Zymo Research, R2062). The RNA fragments corresponding to 17-34 nt in length were excised from the gel, dephosphorylated by T4 polynucleotide kinase (New England Biolabs, M0201S), ligated to linkers by T4 RNA Ligase 2, truncated KQ (New England Biolabs, M0373L), rRNA-depleted by a Ribo-Zero Gold rRNA Removal Kit (Human/Mouse/Rat) (Illumina), and reverse transcribed by ProtoScript II Reverse Transcriptase (New England Biolabs, M0368L). Then, the cDNA was circularized by CircLigase II ssDNA ligase (LGC Biosearch Technologies, CL9025K) and amplified by PCR with Phusion High-Fidelity DNA Polymerase (New England Biolabs, M0530S). DNA libraries were sequenced on HiSeq 4000 (Illumina) with a single-end 50 bp option.

### Library preparation for Thor-Ribo-Seq

For the “aliquoted-then-digested” experiments, cell lysates corresponding to 1, 0.1, and 0.01 µg of RNA were scaled up to 300 µl with lysis buffer, digested with 0.067 U/µl RNase I at 25°C for 45 min, placed on ice, supplemented with 200 U of SUPERase•In RNase Inhibitor, and subjected to a sucrose cushion.

For the “digested-then-aliquoted” experiments, cell lysates were treated with RNase I and SUPERase•In RNase Inhibitor as described in the “*Library preparation for standard Ribo-Seq*”. Then, aliquots corresponding to 1, 0.1, and 0.01 µg of RNA were brought up to a volume of 300 µl with lysis buffer and subjected to a sucrose cushion.

RNA isolation and gel excision were performed as described previously ^8^. The first linker (SI181) 5′-/Phos/NNNNN**ATCGT**AGATCGGAAGAGCACACGTCTGAACTCCAGTCA*CCCT ATAGTGAGTCGTATTA*GTCA/ddC/-3′, where /Phos/ represents a 5′ phosphate, /ddC/ represents a terminal 2′, 3′-dideoxycytidine, N represents a random nucleotide for UMI, bold characters represent a sample barcode sequence, and italicized characters correspond to the T7 promoter, and the second linker SI183 5′-/Phos/NNAGATCGGAAGAGCGTCGTGTAGGGAAAGAG/ddC/-3′ was preadenylated by Mth RNA Ligase (New England Biolabs, E2610S) as described previously ^8^. After dephosphorylation by T4 polynucleotide kinase, RNAs were ligated with the preadenylated first linker SI181 by T4 RNA Ligase 2, truncated KQ. After hybridization of the T7 promoter oligonucleotide SI182 5′-TGAC*TAATACGACTCACTATAGGG*TGACTGGAGTTCAG-3′, complementary RNA was transcribed by a T7-Scribe Standard RNA IVT Kit (CELLSCRIPT) at 37°C for 2.5 h and gel-purified. After dephosphorylation by T4 polynucleotide kinase, RNAs were ligated to the preadenylated second linker SI183 by T4 RNA Ligase 2, truncated KQ, hybridized to SI186 5′-/Phos/AATGATACGGCGACCACCGAGATCTACACTCTTTCCCTACACGACGCTC-3′, and then reverse-transcribed by ProtoScript II Reverse Transcriptase. cDNA was PCR-amplified with NI798 5′-AATGATACGGCGACCACCGAGATCTACACTCTTTCCCTACACGACGCTC-3′ and NI799 5′-CAAGCAGAAGACGGCATACGAGATCGTGATGTGACTGGAGTTCAGACGTGTG-3′ ^7^ by Phusion High-Fidelity DNA Polymerase. rRNA depletion by the Ribo-Zero unit from TruSeq Stranded Total RNA Library Prep Gold was applied to cDNAs, but can be used for the first linker-ligated RNAs. DNA libraries were sequenced on HiSeq X (Illumina) with a paired-end 150-bp option.

### Library preparation for HT-Thor-Ribo-Seq

Briefly, 20 U of RNase I was manually dispensed into each well of a 96-well plate (Eppendorf, 0030129504), followed by the addition of 36 µl of cell lysate. After sealing the plate with aluminum foil (Bio-Rad, 1814045) and briefly mixing, the plate was incubated at 25°C for 45 min and then placed on ice. The lysate in each well was subsequently mixed with 26.6 U of SUPERase•In and transferred to a 0.2-µm filter plate containing 690 µl of Sephacryl S-400 HR (Cytiva, 17060910), which was equilibrated with lysis buffer without Triton X-100. For the optimization experiments shown in Figure S2, the flowthrough was mixed with 10 mM EDTA and subjected to ultrafiltration using AcroPrep Advance 96-well Omega 100K MWCO filter plates (Cytiva, 8036). The samples were then mixed with three volumes of TRIzol LS (Thermo Fisher Scientific, 10296010), incubated at room temperature for 5 min, and subjected to RNA extraction using the Direct-zol 96 RNA kit (Zymo Research, R2056).

The first linkers (SI96_001–096; 5′-/Phos/NNNNN**IIIIIIIIIIIIII**AGATCGGAAGAGCACACGTCTGAACTCCAGTCAC ATTATCCCTATAGTGAGTCGTATTAAATTC/SpC3/, where bold “**I**” denotes a 14-mer barcode listed in Table S5) were pre-adenylated with Mth RNA Ligase, purified using the ZR-96 Oligo Clean & Concentrator (Zymo Research, D4062), and then ligated to dephosphorylated sample RNAs. The 14-mer barcode sequence was referenced from the Quartz-Seq2 protocol ^154^. The samples were then treated with 1 µl of 5′ Deadenylase (New England Biolabs, M0331S) and 1 µl of Lambda Exonuclease (New England Biolabs, M0262L) at 30°C for 45 min, mixed with three volumes of TRIzol LS, and pooled into a single Phasemaker tube (Thermo Fisher Scientific, A33248). The total volume of the pooled TRIzol LS mixture was measured, and 200 µl of chloroform was added for every 750 µl of the pooled mixture, followed by vigorous vortexing and centrifugation at 16,000 × g for 5 min at 4 °C. The aqueous phase was collected and subjected to ethanol precipitation. Purified RNAs were separated by electrophoresis, and the gel regions corresponding to monosomes (17–34 nt) and disomes (50–80 nt) were excised. Subsequent procedures, including rRNA depletion, *in vitro* transcription, second linker ligation, reverse transcription, and PCR amplification, were performed as described in the “*Library preparation for Thor-Ribo-Seq*” section, essentially. To generate the dsDNA T7 promoter for linker-ligated RNA, SI194 5′-GAATTTAATACGACTCACTATAGGGATAATGTGACTGGAGTTCAG-3′ was used. Liquid handling between 96-well plates was automated using an ASSIST PLUS pipetting robot, VOYAGER multichannel pipettes (INTEGRA, 4743, 4721, 4722, and 4724), and a D-ONE pipette (INTEGRA, 4532). DNA libraries were sequenced on HiSeq X (Illumina) with a paired-end 150-bp option.

### Library construction for RNA-Seq

For the siRNA knockdown library, total RNA was extracted from lysates using Trizol LS and Direct-zol 96 RNA kit. After rRNA depletion using Human riboPOOL Standard RNA-Seq (siTOOLs biotech, dp-K096-000054), the library preparation was performed using SEQuoia Express Stranded RNA Library Prep Kit (Bio-Rad, 12017265) and SEQuoia Dual Indexed Primers Plate (Bio-Rad, 12011930). Liquid handling was automated using an ASSIST PLUS pipetting robot and VOYAGER multichannel pipettes. DNA libraries were sequenced on NovaSeq X Plus (Illumina) with a paired-end 150-bp option.

For PBMCs, total RNA was extracted from frozen PBMCs using TRIzol reagent (Thermo Fisher Scientific, 15596018) and a Direct-zol RNA MicroPrep (Zymo Research, R2062). Library construction and sequencing were performed manually using a TruSeq stranded mRNA Library Prep kit (Illumina) and an Illumina NovaSeq6000 with a paired-end 100-bp option or an Illumina NextSeq2000 with a single-end 65-bp option.

### Deep sequencing data analysis

Data processing was performed as previously described ^7^ with modifications. Using Fastp v0.21.0 ^155^, bases of pair-end reads were corrected by overlap analysis, and then quality filtering and removal of the adapter sequence (AGATCGGAAGAGCACACGTCTGAA) were performed on read 1. For single-end reads of RNA-Seq, base correction was skipped. After the demultiplexing by barcode sequences on the linkers using cutadapt v3.7 ^156^ and the extraction of UMI by UMItools v1.1.2 ^157^, reads mapped to noncoding RNAs using STAR v2.7.0a ^158^ were excluded from the analysis. The remaining reads were aligned to the human genome hg38 and assigned to the GENCODE Human release 44 reference using STAR. UMI suppression was performed using UMItools. Downsampling of datasets was performed using seqtk (https://github.com/lh3/seqtk).

The offsets of the A site were determined based on the metagene analysis for the 5′ end of footprints (Table S6). For RNA-Seq, offsets were set to 15 for all mRNA fragments. The periodicity scores, defined as the product of the relative entropy and the fraction of each footprint length, were calculated as described in ^42^.

For ORF-wise analysis, the canonical transcripts were defined based on MANE Select v1.0 ^159^, supplemented with the Ensembl Canonical for genes not included in the MANE Select list. Reads for each gene were counted after excluding the first and last five codons of ORFs. The fold changes of footprints, RNAs, and translation efficiencies, which are footprints normalized by RNA-Seq counts, were calculated using the DESeq2 package ^160^ with the likelihood ratio test.

GO terms and domain information of genes were obtained from the Uniprot database (https://www.uniprot.org/, on February 27th, 2025). Similar terms were manually aggregated. GO analysis for COVID-19 patients was performed with DAVID (https://david.ncifcrf.gov/home.jsp) ^161,162^. Dendrograms for changes of ribosome footprints, RNA, and translation efficiency were depicted using the ggtree package ^163^, with hierarchical clustering performed using Spearman’s rank correlation coefficient and Ward.D2 linkage. Partitioning Around Medoids (PAM) clustering was performed using the cluster package ^164^. Periodicity of footprints was calculated by the periodogram function in the GeneCycle package ^165^. The RG4 annotation was based on all genes in the list of the union RG4s across rG4-Seq 2.0 libraries ^90^.

Ribosome occupancies on each codon were defined as the number of reads at given codons normalized by the average reads per codon on each ORF. Transcripts with more than 1 (for standard and Thor-Ribo-Seq shown in Figure 1) or 0.1 (for HT-Thor-Ribo-Seq) reads per codon on average in all samples were considered.

### ENCODE eCLIP-Seq data analysis

The BAM files of eCLIP-Seq data aligned to hg38 were obtained from the ENCODE portal (https://www.encodeproject.org). The 5′ ends of reads corresponding to each region (5′ UTR, ORF, and 3′ UTR) were counted. The fold change of IP samples relative to size-matched input samples in each region was then calculated using DESeq2. Only genes with a read count of at least one were included in the analysis.

### On-gel mito-FUNCAT

On-gel mito-FUNCAT was conducted as described previously ^119^. HEK293 cells were transfected with 5 μM siRNA targeting IGF2BP3 (Horizon Discovery, L-003976-00-0005) or control siRNA (Horizon Discovery, D-001206-13-20) using *Trans*IT-X2 in a 6-well plate and incubated for 3 d. Then, cells were cultured in methionine-free medium supplemented with 50 µM HPG (Jena Bioscience, CLK-1067) and 100 µg/ml anisomycin (Alomone Labs, A520) for 3 h before the cells were harvested. Following cell lysis, nascent peptides were labeled with 50 μM IRDye800CW Azide (LI-COR Biosciences, 929-60000) using the Click-iT Cell Reaction Buffer Kit (Thermo Fisher Scientific, C10269) following the manufacturer’s protocol. The reactants were separated on SDS– PAGE gels, and signals were captured using the Odyssey CLx imaging system (LI-COR Biosciences) via the IR 800-nm channel. Signal intensities were normalized based on Coomassie Brilliant Blue (CBB) staining (GelCode Blue Safe Protein Stain, Thermo Fisher Scientific, 24594), which was visualized using the IR 700-nm channel in an Odyssey CLx. Image quantification was performed with Image Studio software (LI-COR Biosciences, v5.2).

### Construction of RQC reporter plasmids

To construct the tandem ORF reporter plasmid, DNA fragments containing EGFP-P2A-FLAG and P2A-HA-DsRedEx were PCR-amplified. These two fragments were joined by PCR and cloned into pCS2+MT containing the zebrafish suv39 h1a 3′ UTR ^166^ using the NcoI and XhoI sites. To insert the Lys (AAA) ×15 sequence, oligonucleotides were annealed and cloned downstream of the FLAG tag using the EcoRV and EcoRI sites.

### FACS

Cells seeded in a 6-well plate were transfected with 5 μM siRNA targeting RACK1 (Horizon Discovery, L-006876-00-0005), HUWE1 (Horizon Discovery, L-007185-00-0005), NKRF (Horizon Discovery, L-031862-01-0005), or control siRNA (Horizon Discovery, D-001810-10-50) using *Trans*IT-X2 and incubated for 24 h. The reporter plasmids were then introduced using *Trans*IT-X2 reagent and incubated for 48 h. Pre-treatment for FACS was performed as described previously ^167^. The cells were trypsinized (Thermo Fisher Scientific, 25300-54) and analyzed via a FACSAria II Special Order system (BD) with FACSDiva v6.1.3 software (BD). The gating strategy can be found in Figure S5H.

### Statistical analysis and reproducibility

All statistical analysis was done using R v4.3.0 on R Studio (v2024.12.0+467). The significance of the differential expression analysis was calculated by a likelihood ratio test in a generalized linear model using the DESeq2 package. The two-tailed Mann– Whitney *U* test with Benjamini-Hochberg adjustment was used for Figures 3C-E, 5D, 6B, and S3K-L. For Figures 5E–F, 7B, and S3M, the Mann–Whitney *U* test was performed without multiple testing correction. The two-sided Tukey-Kramer multiple comparison test was used for comparisons of the mean value in Figures 5I, S5G, and S7P. The two-sided Student’s t-test was used for comparisons in Figures 6C and S5I. For western blotting, FACS, and Mito-FUNCAT, we performed the experiments independently three times.

## Data availability

Standard Ribo-Seq, Thor-Ribo-Seq data (GEO: GSE222195), and HT-Thor-Ribo-Seq/RNA-Seq (GEO: GSE304445) were deposited in the National Center for Biotechnology Information (NCBI) database. The previous disome profiling data (GEO: GSE145723) were used.

## Code availability

The custom scripts used in this study are available at Zenodo (DOI: 10.5281/zenodo.16730947).

---

**Figure S1.**
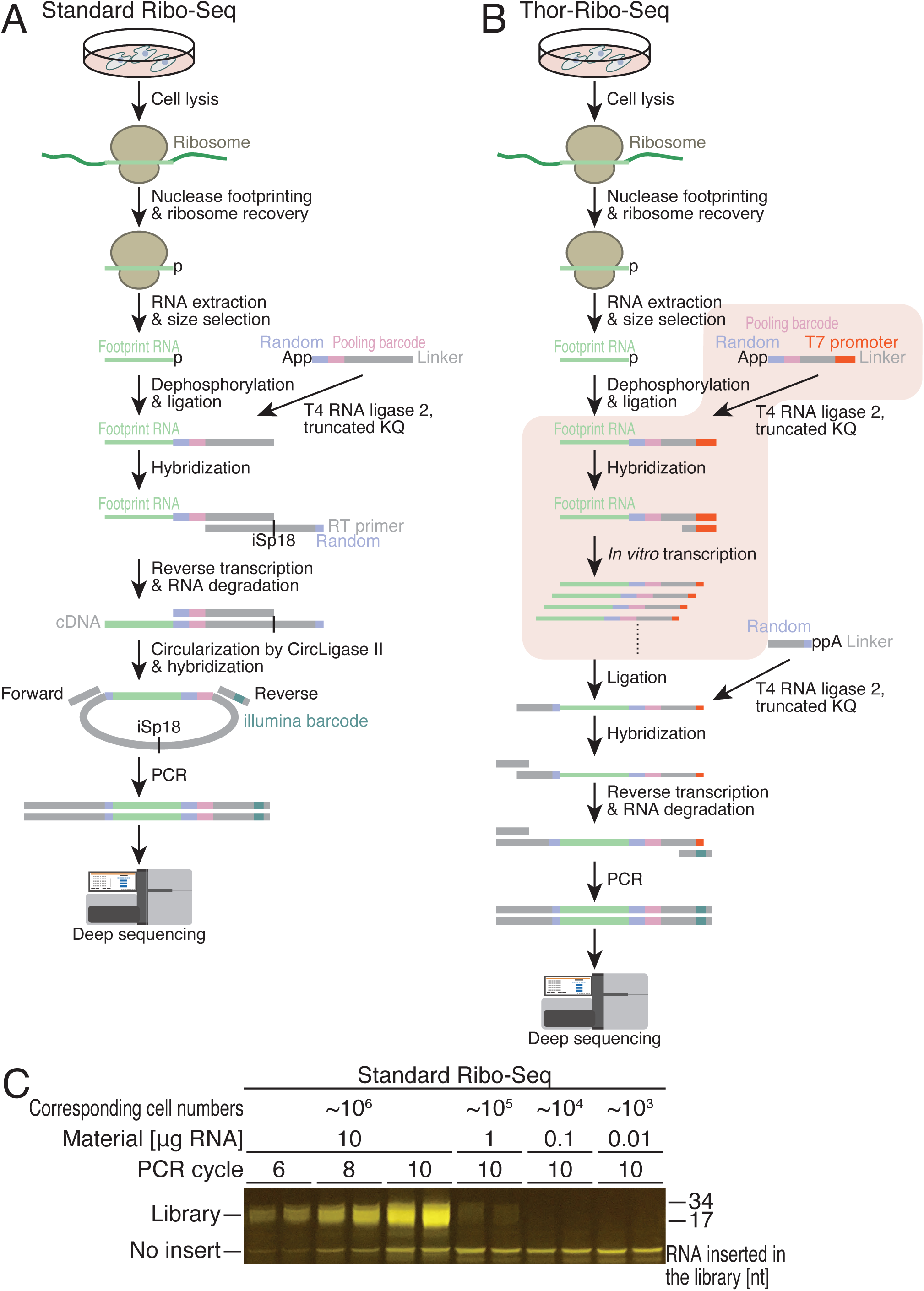
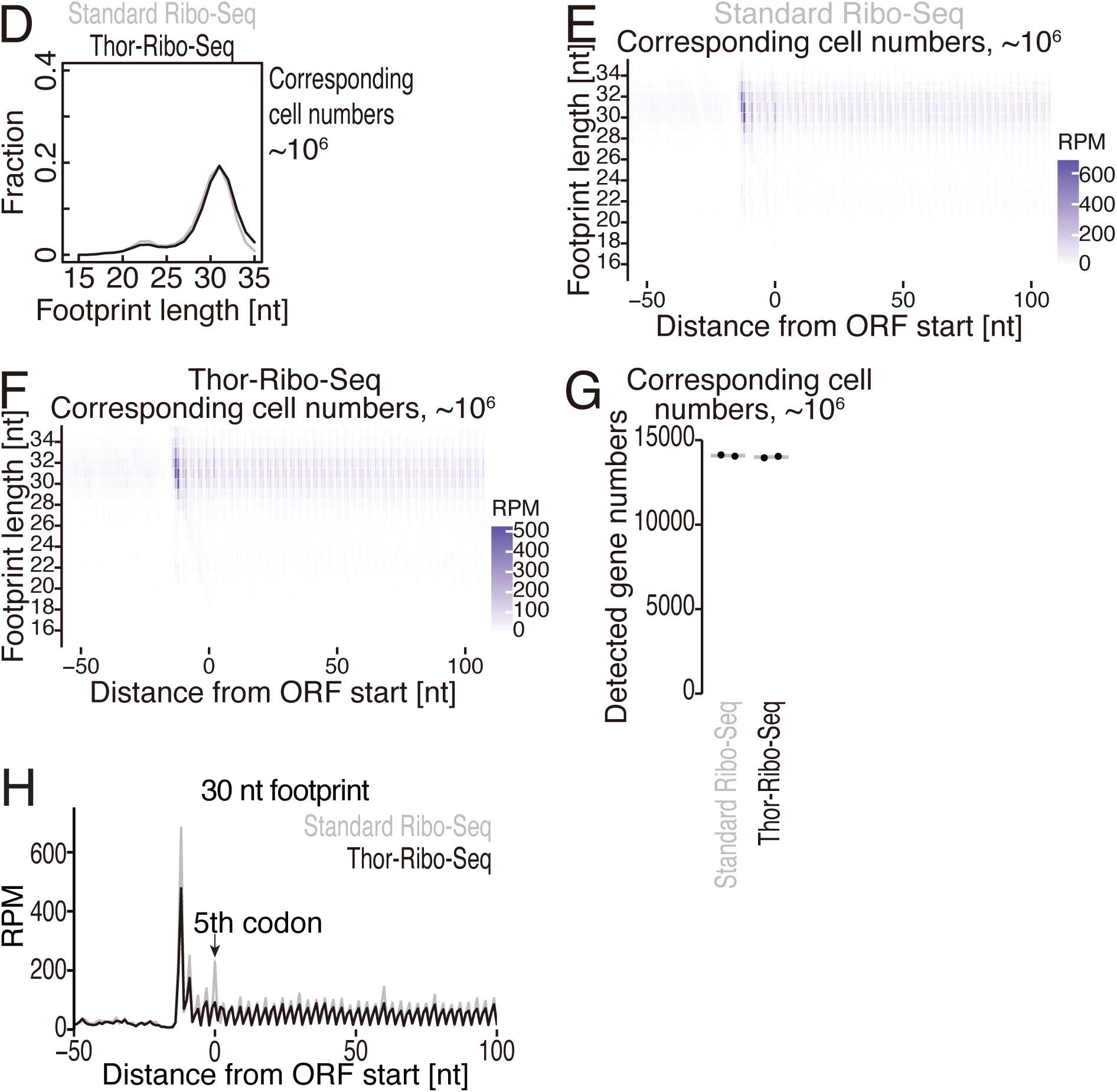
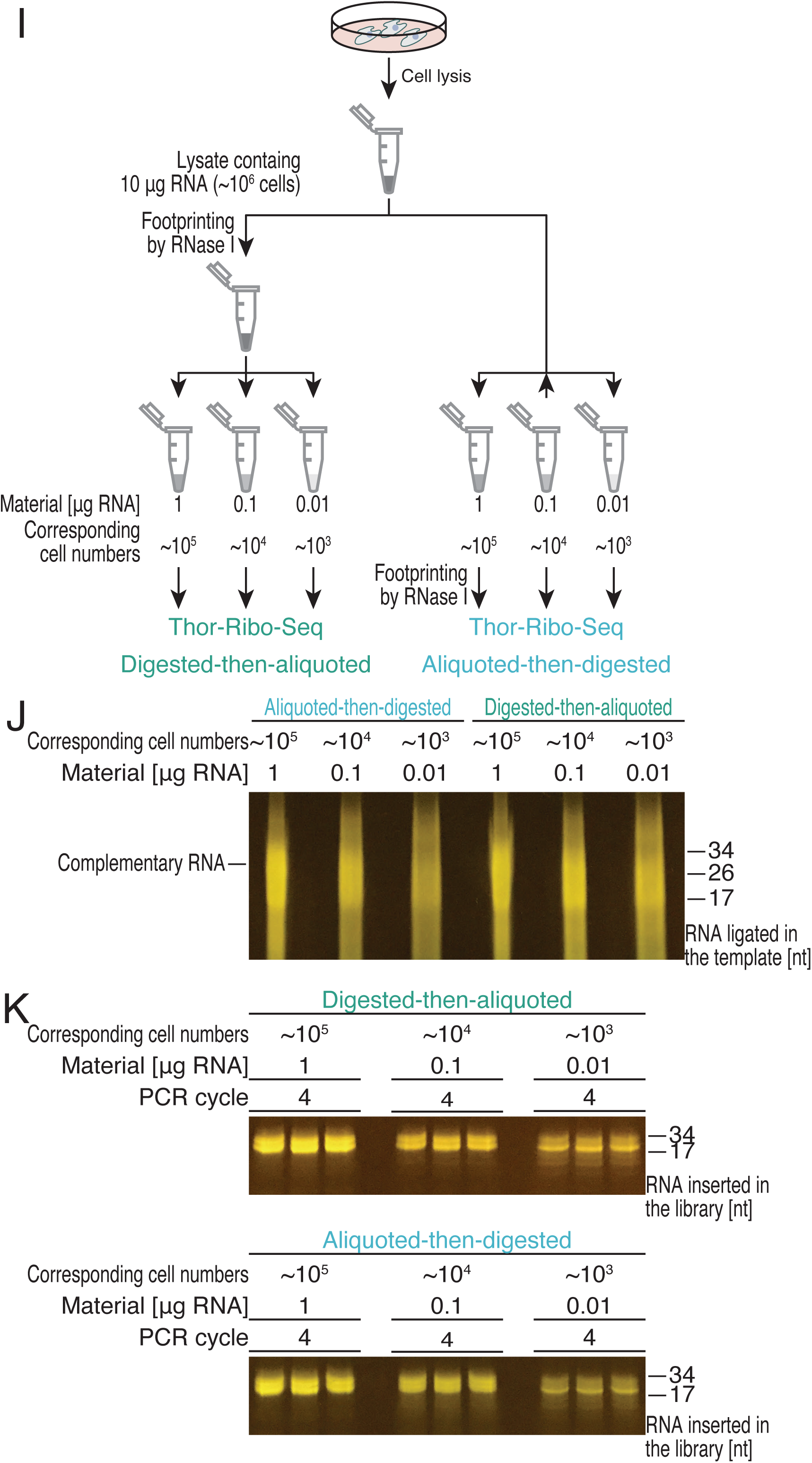
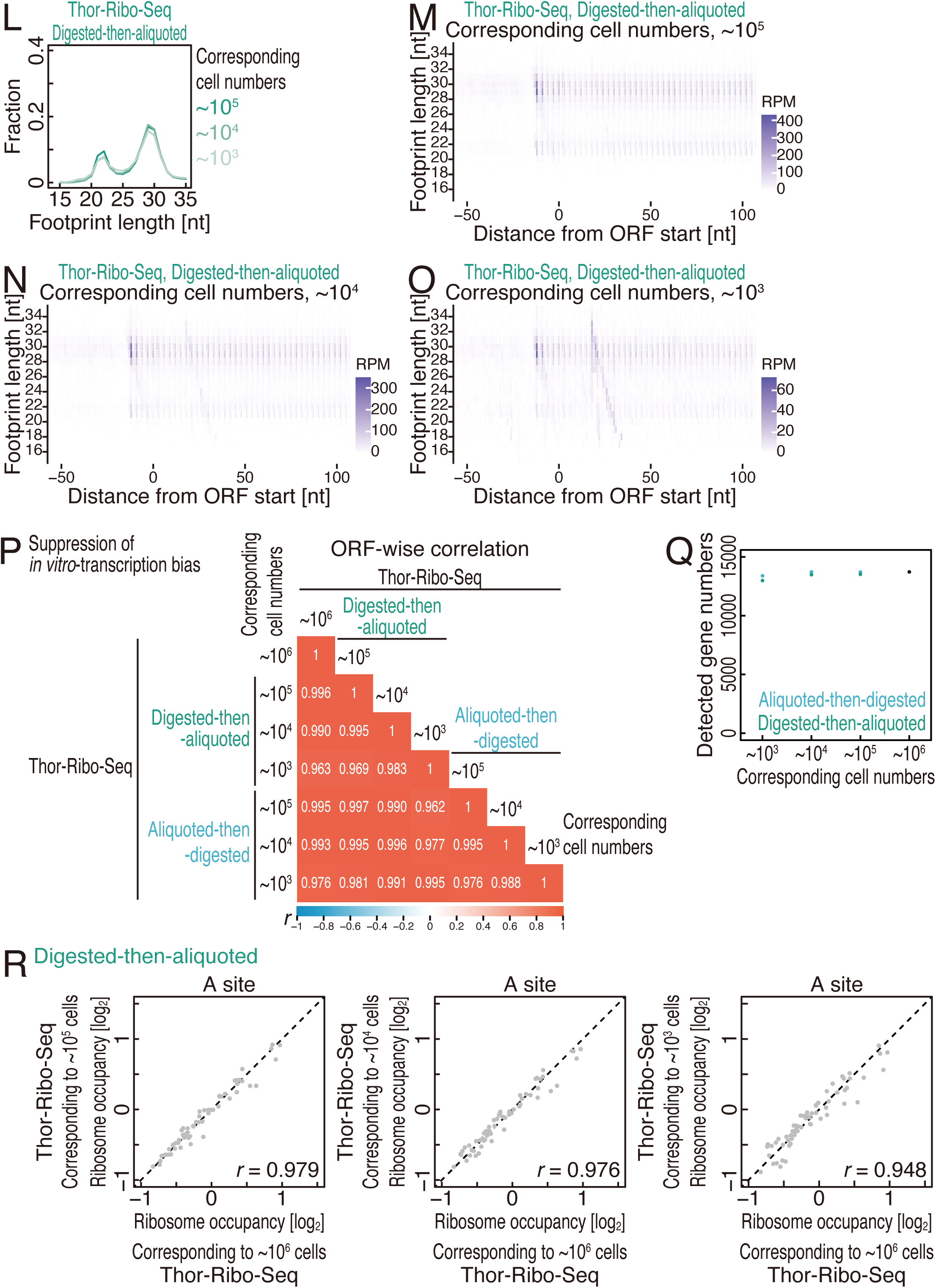
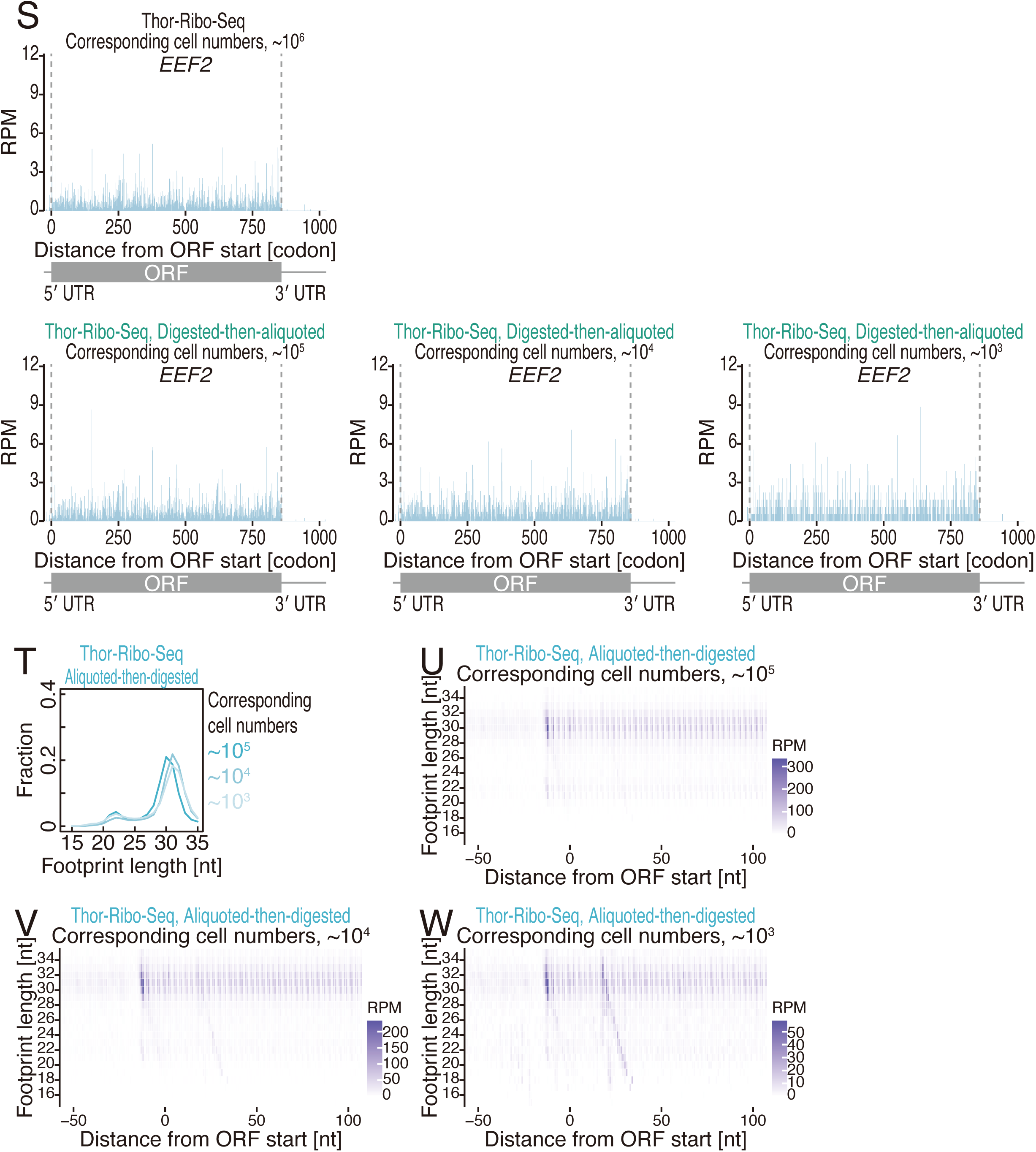
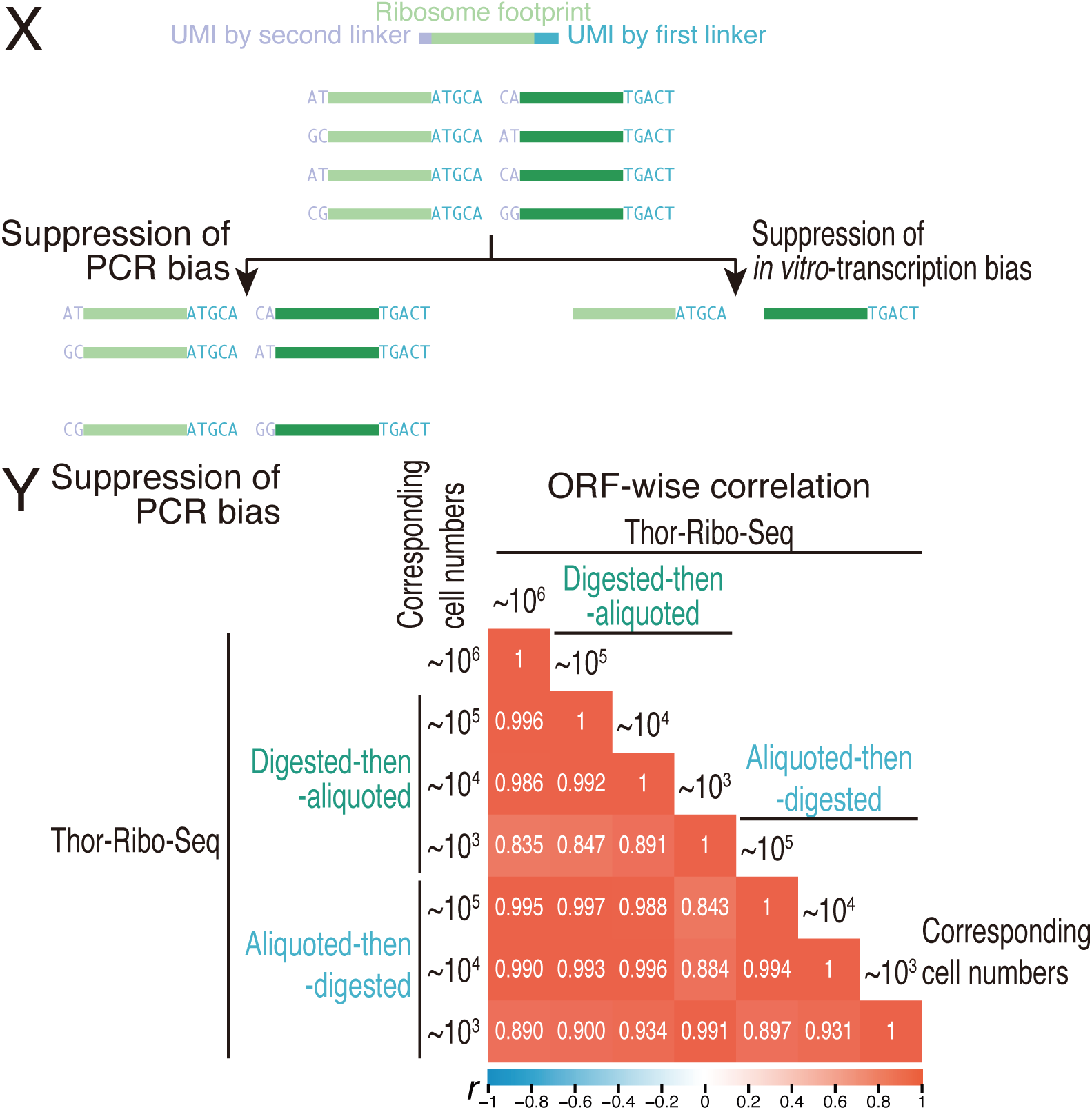
Library preparation and investigation for biases in Thor-Ribo-Seq. **Related to Figure 1**. (A and B) Step-by-step overviews of library preparation schemes for standard Ribo-Seq (A) and Thor-Ribo-Seq (B). Random sequences were used as UMI. (B) A gel image of the PCR-amplified DNA library prepared by the standard Ribo-Seq method with titrated amounts of initial materials. (C) The distribution of footprint length for standard Ribo-Seq and Thor-Ribo-Seq with normal inputs. (E and F) Metagene plots of the 5′ end position of ribosome footprints around start codons in the indicated experiments. X axis, the position relative to the start codon (0 as the first nucleotide of the start codon); Y axis, footprint length; color scale, read abundance. (G) Detected gene numbers (1 read or more) in 3 million footprint reads in the indicated experiments. (H) Metagene plots of the 5′ end position of 30-nt ribosome footprints around start codons in the indicated experiments. (I) Schematic representation of experimental designs to test the potential of Thor-Ribo-Seq. In the “digested-then-aliquoted” experiments, lysates were treated with RNase I and then divided into different amounts. In the “aliquoted-then-digested” experiments, lysates were first titrated and then digested with RNase I. (J) A gel image of *in vitro* transcription-amplified complementary RNA to the ribosome footprints. (K) A gel image of the PCR-amplified DNA library prepared by the Thor-Ribo-Seq method with titrated amounts of initial materials. (L and T) The distribution of footprint length for Thor-Ribo-Seq with “digested-then-aliquoted” conditions (L) and that with “aliquoted-then-digested” conditions (T). (M-O and U-W) Metagene plots of the 5′ end position of ribosome footprints around start codons in the indicated experiments. X axis, the position relative to the start codon (0 as the first nucleotide of the start codon); Y axis, footprint length; color scale, read abundance. (P) Pearson’s correlation coefficient (*r*) among the indicated experiments. Read duplications generated by *in vitro* transcription were suppressed by the UMI in the first linker. The *r* value scales are shown at the color bars. (Q) Detected gene numbers (1 read or more) in 4 million footprint reads in the indicated experiments. (R) Correspondence of averaged ribosome occupancy on A-site codon sequences across Thor-Ribo-Seq (digested-then-aliquoted) with low material inputs. *r*, Pearson’s correlation coefficient. (S) Ribosome footprint distribution on *EEF2* mRNA of Thor-Ribo-Seq with “digested-then-aliquoted” experiments with low material inputs. (X) Schematic representation of UMI-mediated duplicated read suppression. (Y) Pearson’s correlation coefficient (*r*) among the indicated experiments. Read duplications generated by PCR were suppressed by the UMI in the first and the second linkers. The *r* value scales are shown at the color bars. RPM, reads per million mapped reads.

**Figure S2.**
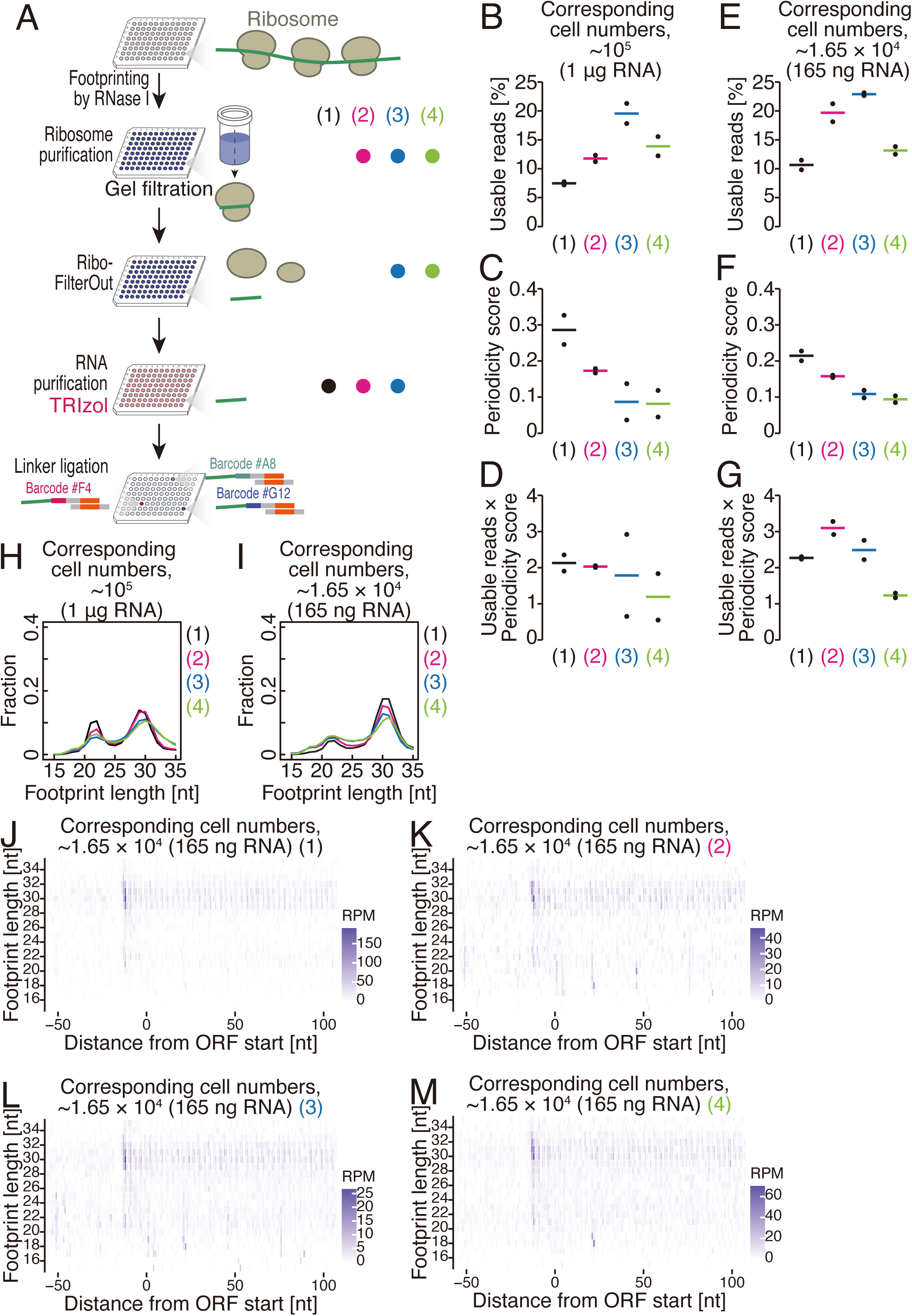
Optimization for HT-Thor-Ribo-Seq. **Related to Figure 2**. (A) Schematic representation of optimization for HT-Thor-Ribo-Seq after the footprinting. Four experimental conditions, each representing a different combination of gel filtration, Ribo-FilterOut, and RNA purification, were tested. (B and E) The ratio of usable reads (genome-mapped reads unmatched to non-coding RNAs such as rRNAs after UMI deduplication) for lysates containing 1 µg (B) or 165 ng (E) RNA, under four conditions. (C and F) Periodicity score for lysates containing 1 µg (C) or 165 ng (F) RNA, under four conditions. (D and G) Values obtained by multiplying the ratio of usable reads by the periodicity score for lysates containing 1 µg (D) or 165 ng (G) RNA, under four conditions. (H and I) The distribution of footprint length for lysates containing 1 µg (H) or 165 ng (I) RNA, under four conditions. (J-M) Metagene plots of the 5′ end position of ribosome footprints around start codons for lysates containing 165 ng RNA, under four conditions. X axis, the position relative to the start codon (0 as the first nucleotide of the start codon); Y axis, footprint length; color scale, read abundance. RPM, reads per million mapped reads.

**Figure S3.**
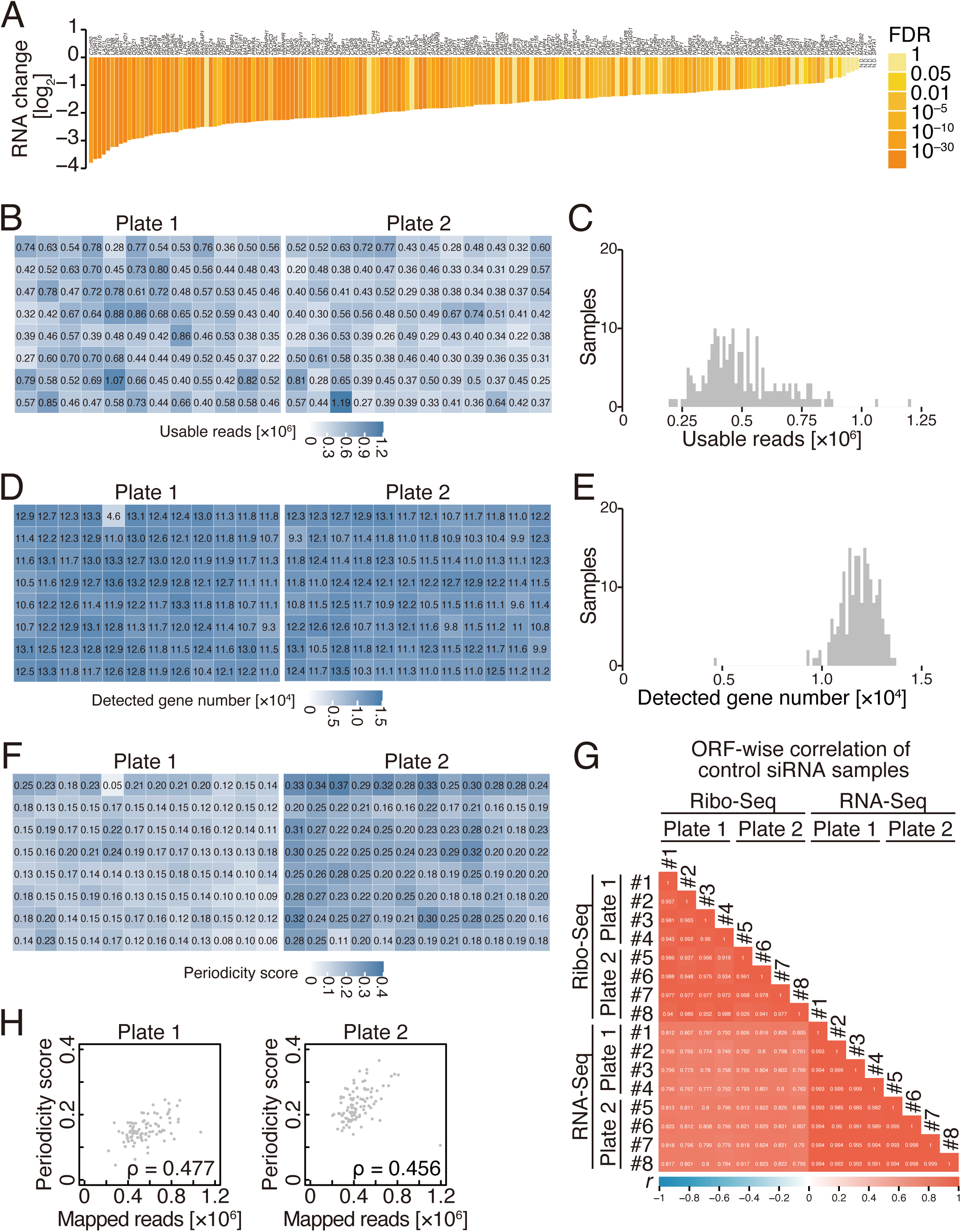
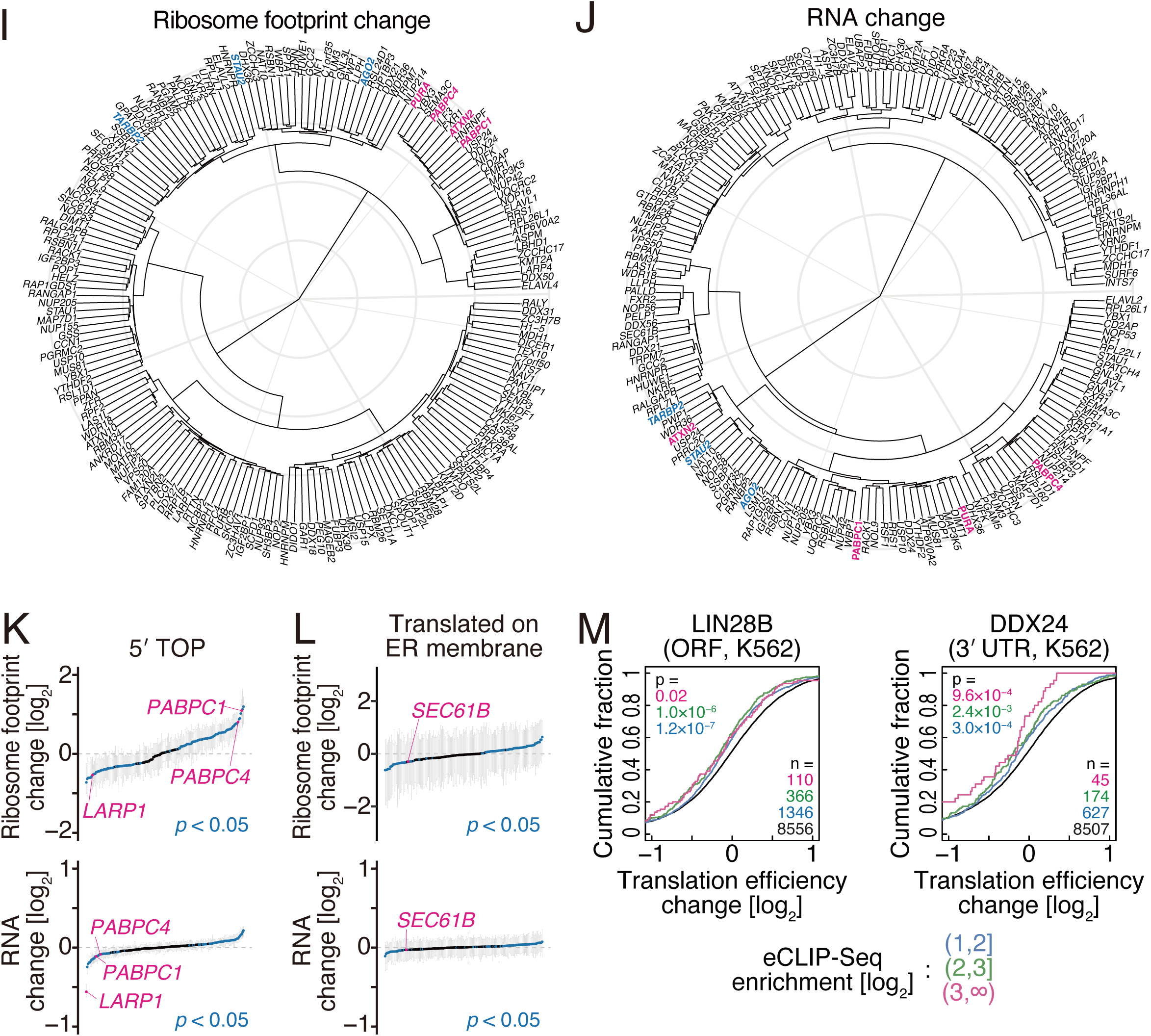
Translation landscape revealed by HT-Thor-Ribo-Seq for 192 siRNA knockdown samples. **Related to Figures 2 and 3**. (A) RNA changes of siRNA-targeted genes after the knockdown in 179 samples. Color represents the false discovery rate (FDR). N.D., not detected. (B, D, and F) Plate view of the number of usable reads (B), detected gene numbers (1 read or more) (D), and periodicity scores (F) for 192 samples. Color scales indicate each value. (C and E) Histogram of the number of usable reads (C) and detected genes (E) for 192 samples. (G) Pearson’s correlation coefficient (*r*) among the control knockdown samples in HT-Thor-Ribo-Seq and RNA-Seq. The *r* value scales are shown at the color bars. (H) Scatter plot of usable reads versus periodicity scores. ρ, Spearman’s rank correlation coefficient. (I and J) Dendrogram of ribosome footprint (I) and RNA (J) changes in 179 siRNA knockdown samples, relative to control samples. Knocked-down genes mentioned in the main text are highlighted. (K and L) Ribosome footprint and RNA changes of mRNAs with a 5′ TOP motif (K) and translated on the ER membrane (L) in 179 siRNA knockdown samples. The median (dot) and first and third quartiles (ticks) are shown. Knockdown samples showing significant up- or down-regulation of indicated mRNAs (*p* < 0.05, two-tailed Mann–Whitney *U* test with Benjamini-Hochberg adjustment) and samples mentioned in the main text are highlighted in blue and magenta, respectively. (M) Cumulative distributions of translation efficiency changes in the knockdown of indicated genes. Genes with log_2_(eCLIP-Seq enrichment) in the intervals (1, 2], (2, 3], and (3, ∞) are shown in blue, green, and magenta, respectively. The *p* values were calculated with the two-tailed Mann‒Whitney *U* test. The mRNA number (n) of each group is shown.

**Figure S4.**
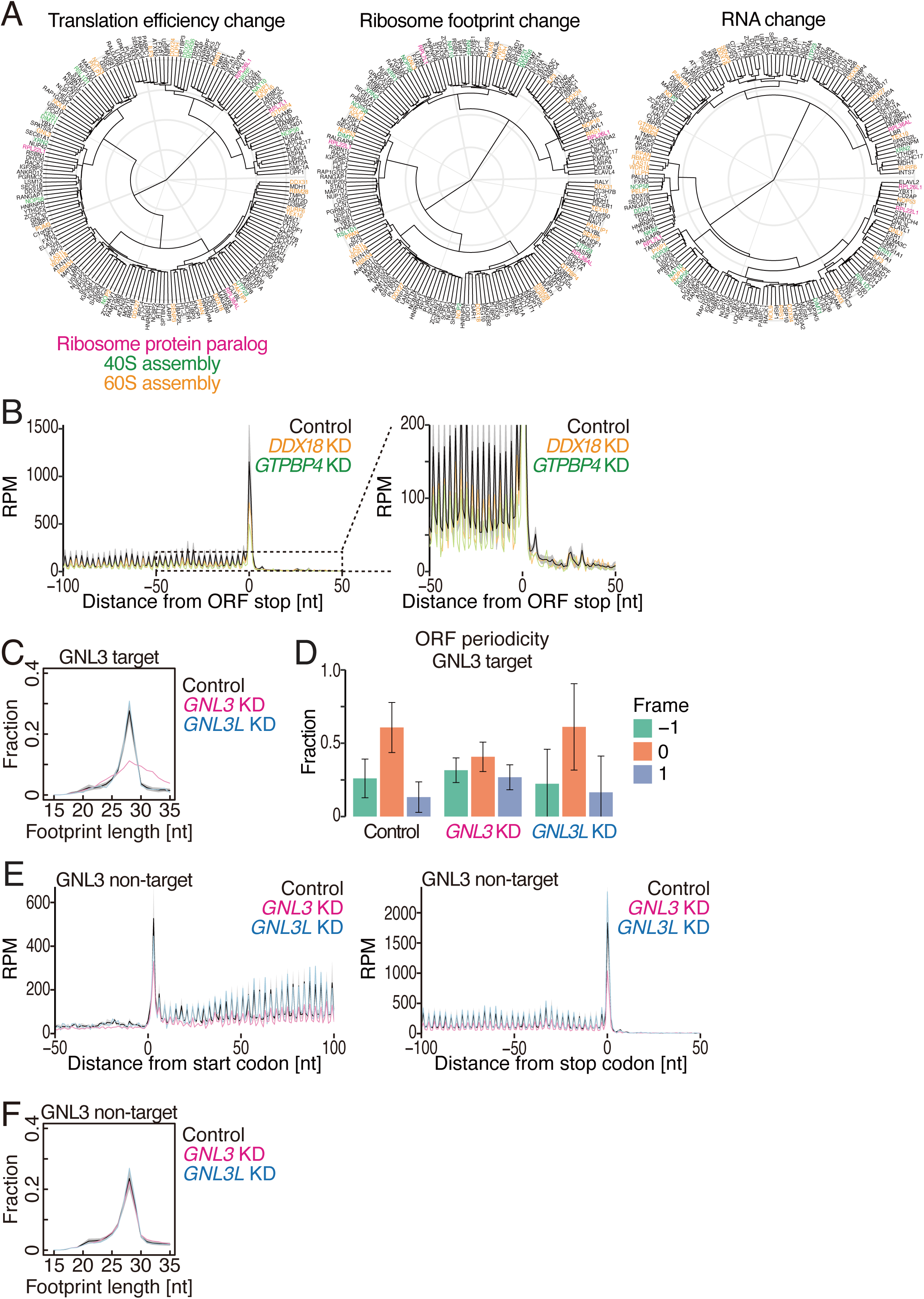
The effect of GNL3 on ribosome movement in UTRs. **Related to Figure 4**. (A) Same as Figures 3B and S3I-J, but genes encoding ribosome protein paralogs (magenta), 40S assembly (green), and 60S assembly (orange) are highlighted. (B) Metagene plots of the 5′ end position of ribosome footprints around stop codons in the indicated knockdown samples. An expanded view is shown on the right. For control knockdown samples, the average across 8 samples is shown as a black line, and the shaded area represents ±1 standard deviation (SD). (C and F) Distribution of footprint lengths for GNL3 target mRNAs (defined in Figure 4D) (C) and GNL non-target mRNAs (F) in the indicated knockdown samples. For control knockdown samples, the average across 8 samples is shown as a black line, and the shaded area represents ±1 standard deviation (SD). (D) Frame distribution of the A-site positions of ribosome footprints within the ORF region for GNL3 target mRNAs (defined in Figure 4D). The means (bars) and SDs (errors) across GNL3 target mRNAs are shown. For control knockdown, values are based on the merged results from 8 samples. (E) Metagene plots for GNL3 non-target mRNAs showing the 5′ end positions of ribosome footprints around start and stop codons. For control knockdown samples, the average across 8 samples is shown as a black line, and the shaded area represents ±1 standard deviation (SD).

**Figure S5.**
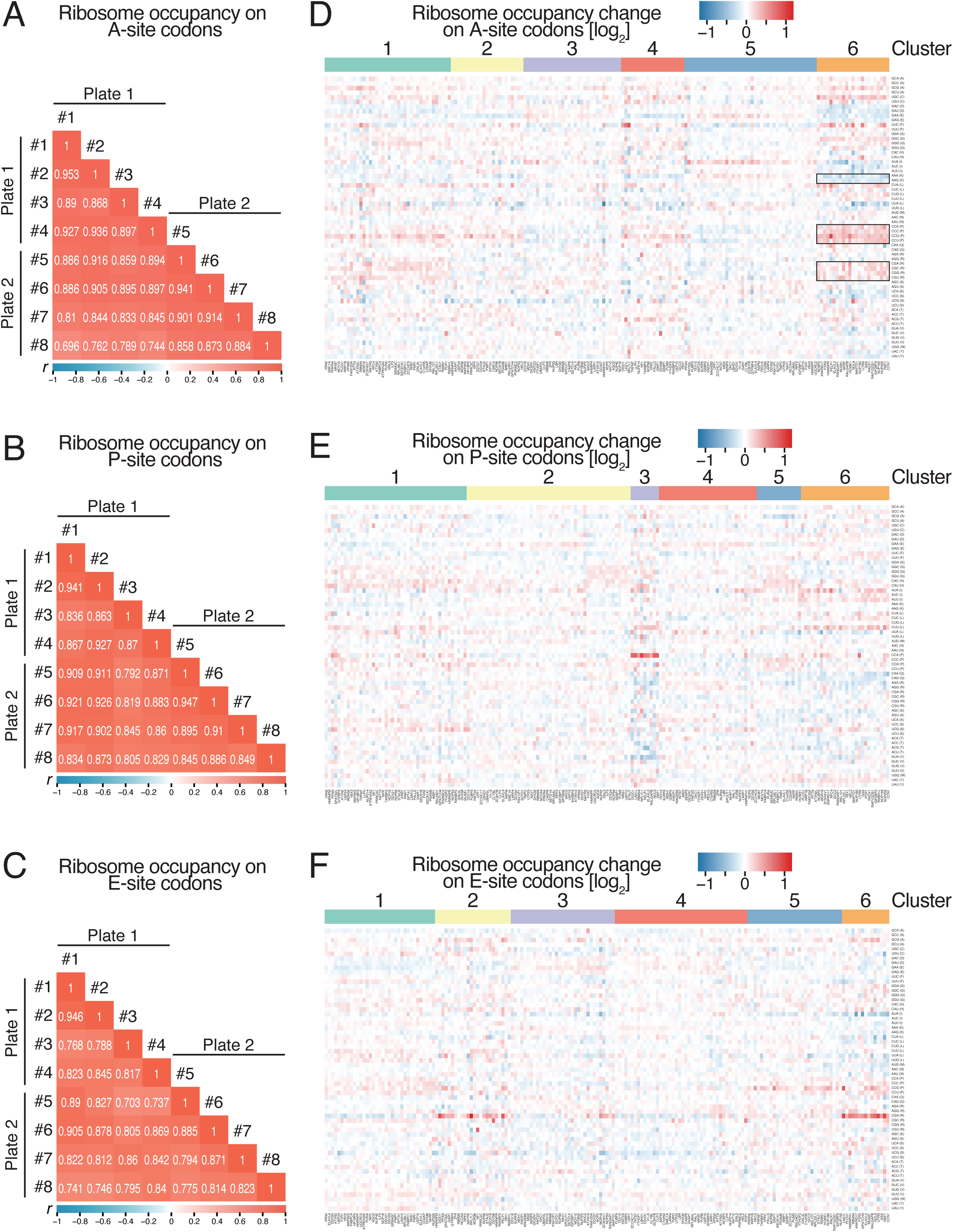
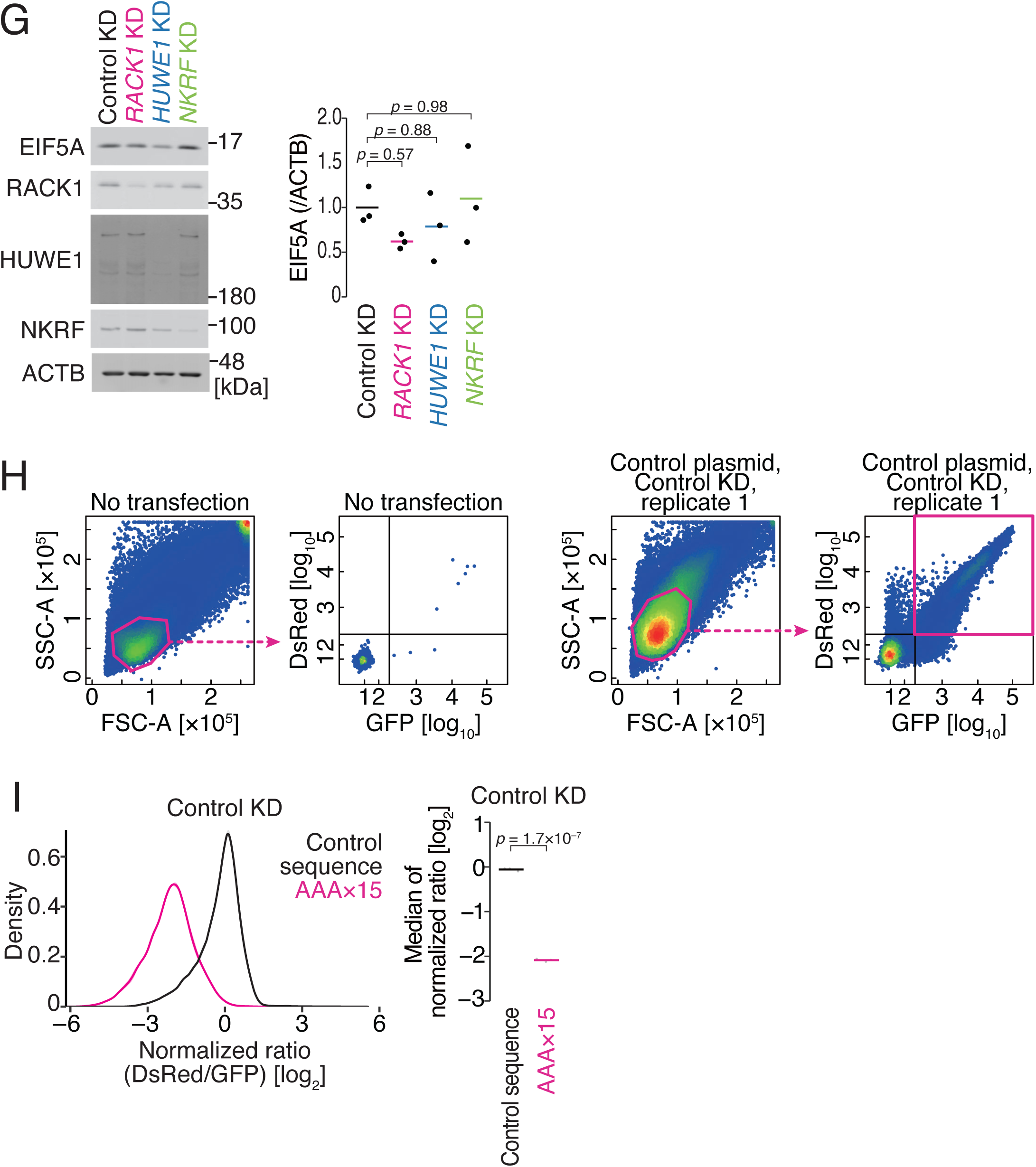
Codon-wise ribosome occupancies revealed by HT-Thor-Ribo-Seq. **Related to Figure 4**. (A-C) Pearson’s correlation coefficient (*r*) of codon-wise ribosome occupancies on A site (A), P site (B), and E site (C) among the control knockdown samples in HT-Thor-Ribo-Seq. The *r* value scales are shown at the color bars. (D-F) Heatmap of codon-wise ribosome occupancy changes on A site (D), P site (E), and E site (F) compared to control samples. The color scale indicates the degree of fold change. The result of partitioning around medoids (PAM) clustering is shown. Codons mentioned in the main text are highlighted in black boxes. (G) Western blotting of the indicated proteins. Quantification of band intensities is shown on the right. The *p* values were calculated by the two-sided Tukey-Kramer multiple comparison test. (H) Representative gating strategy. After excluding doublets and debris, GFP- and DsRed-positive cells were selected. This strategy was applied to all FACS experiments in this study. (I) Density plot (left) and median values (right) of the normalized ratio between DsRed and GFP signals for the indicated reporters in the control knockdown. The *p* value was calculated by the two-sided Student’s t-test. KD, knockdown.

**Figure S6.**
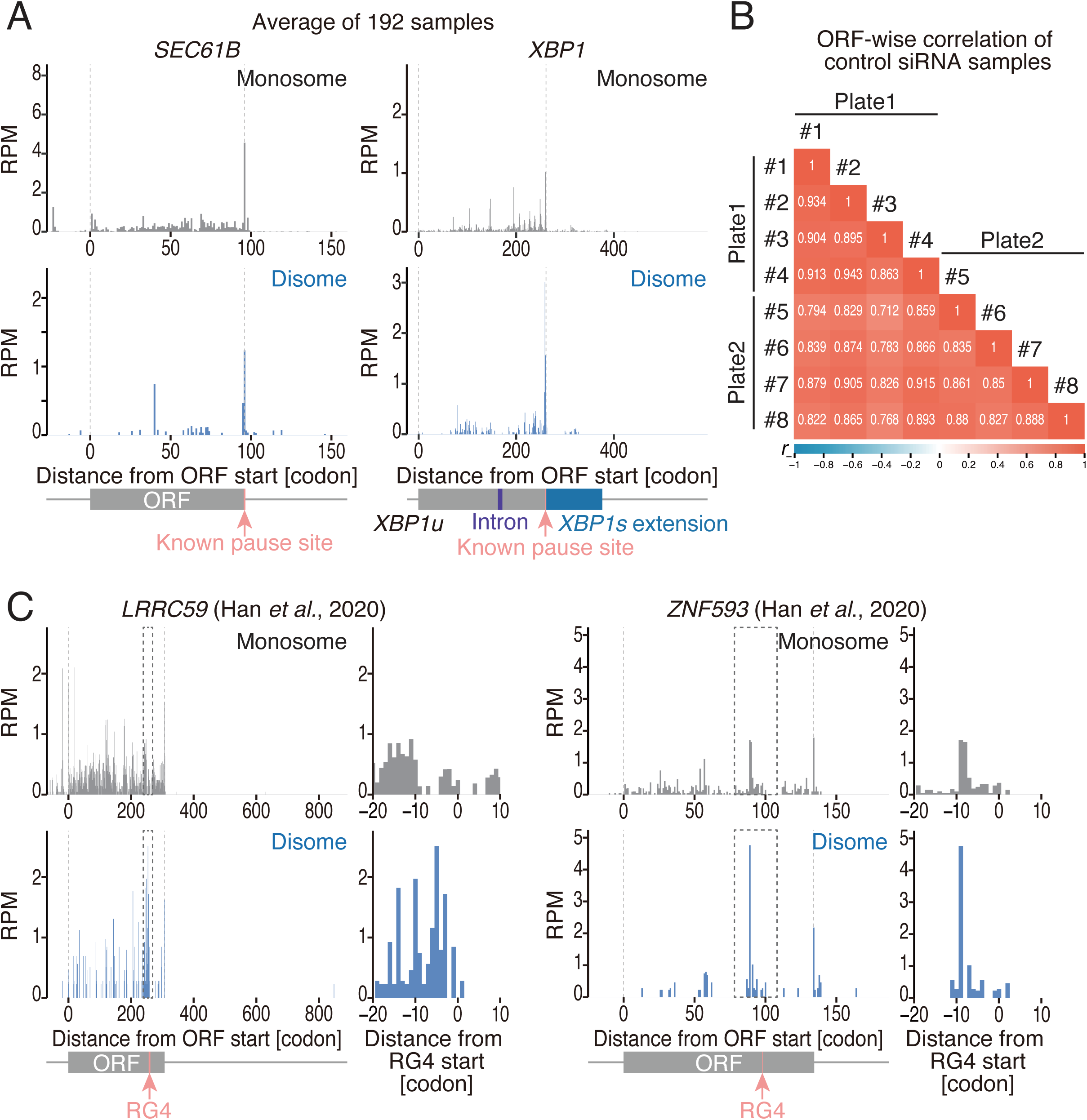
Ribosome collisions revealed by HT-Thor-Disome-Seq. **Related to Figure 5**. (A) Monosome and disome footprint distribution on *SEC61B* and *XBP1* mRNAs. The average across 192 samples is shown. The known ribosome pause sites are shown as pink arrows. (B) Pearson’s correlation coefficient (*r*) of ORF-wise disome footprint counts among the control knockdown samples. The *r* value scales are shown at the color bars. (C) Monosome and disome footprint distribution on indicated RG4-containing mRNAs. Monosome and disome profiling data from ^30^ were analyzed. An expanded view of the dashed line region (around RG4 start position) is shown on the right. RG4 sites are shown as pink arrows. RPM, reads per million mapped reads.

**Figure S7.**
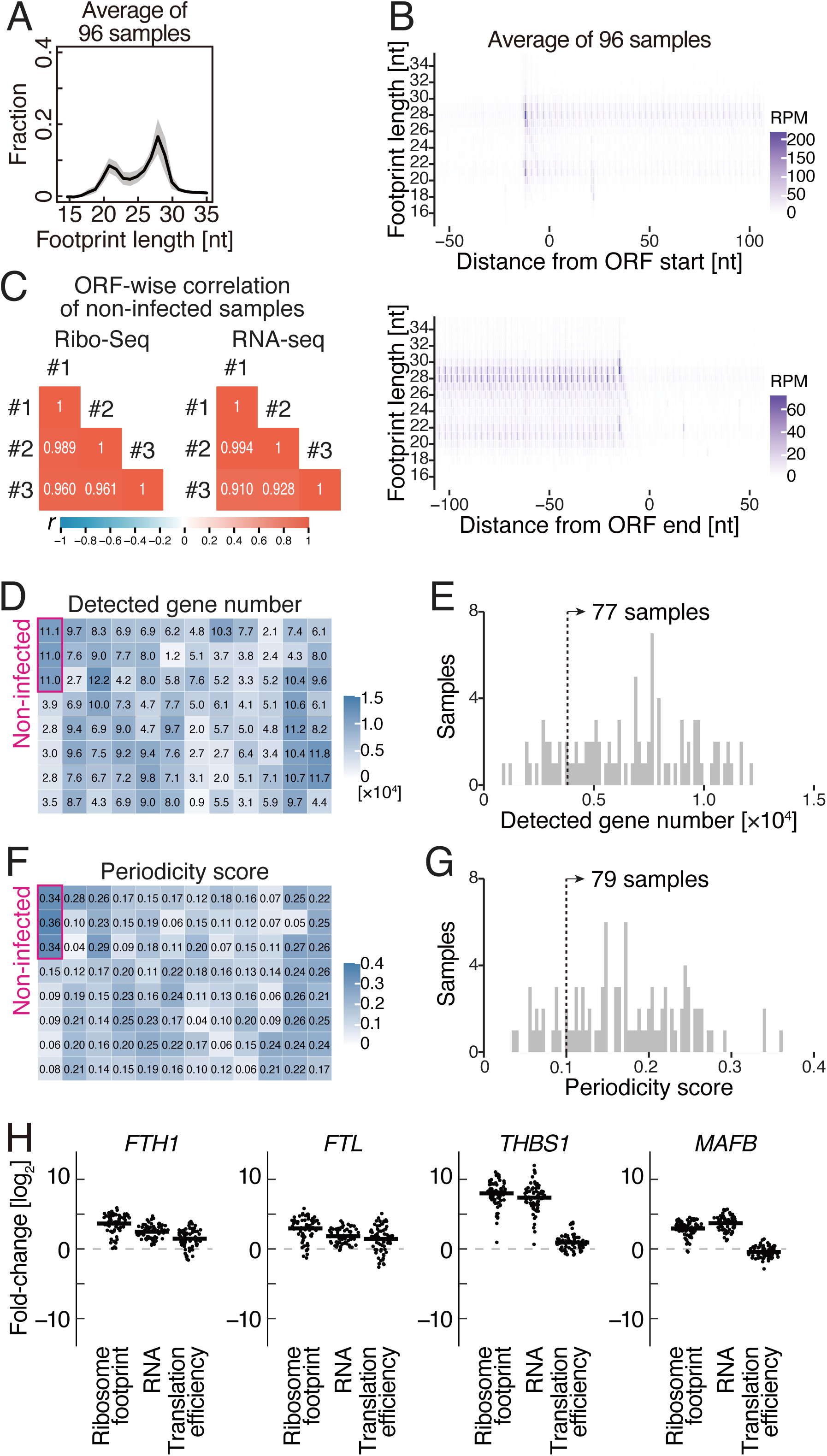
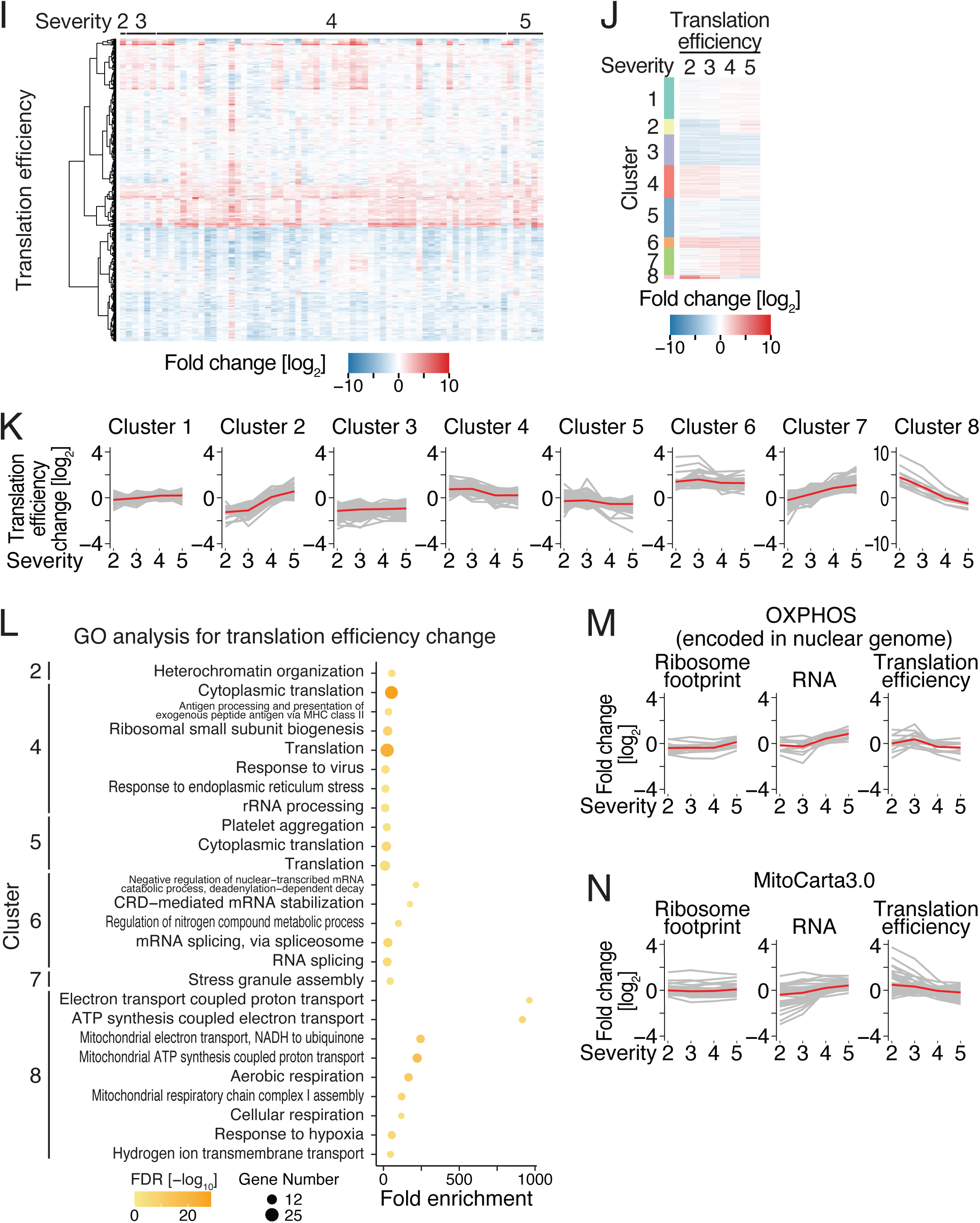
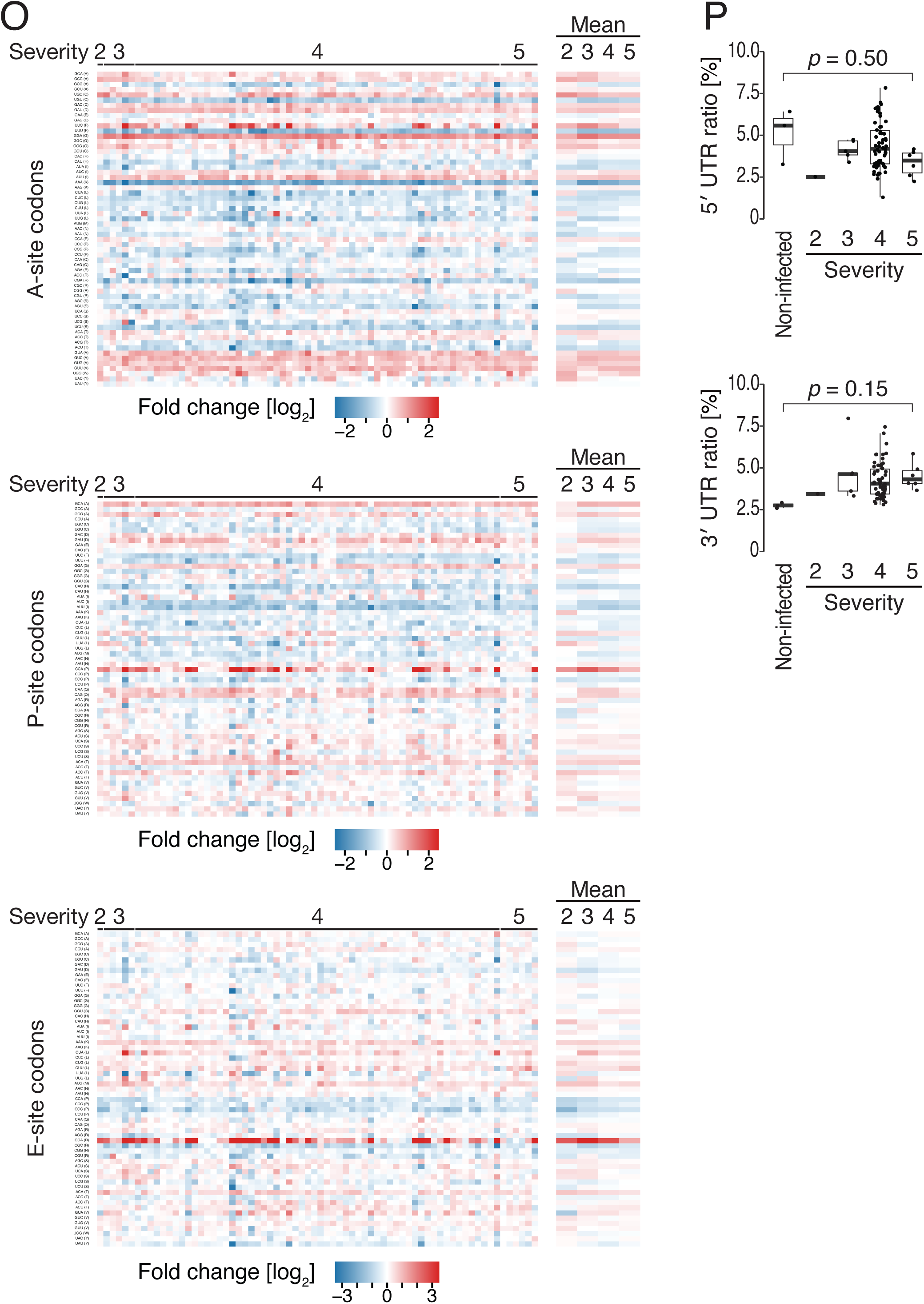
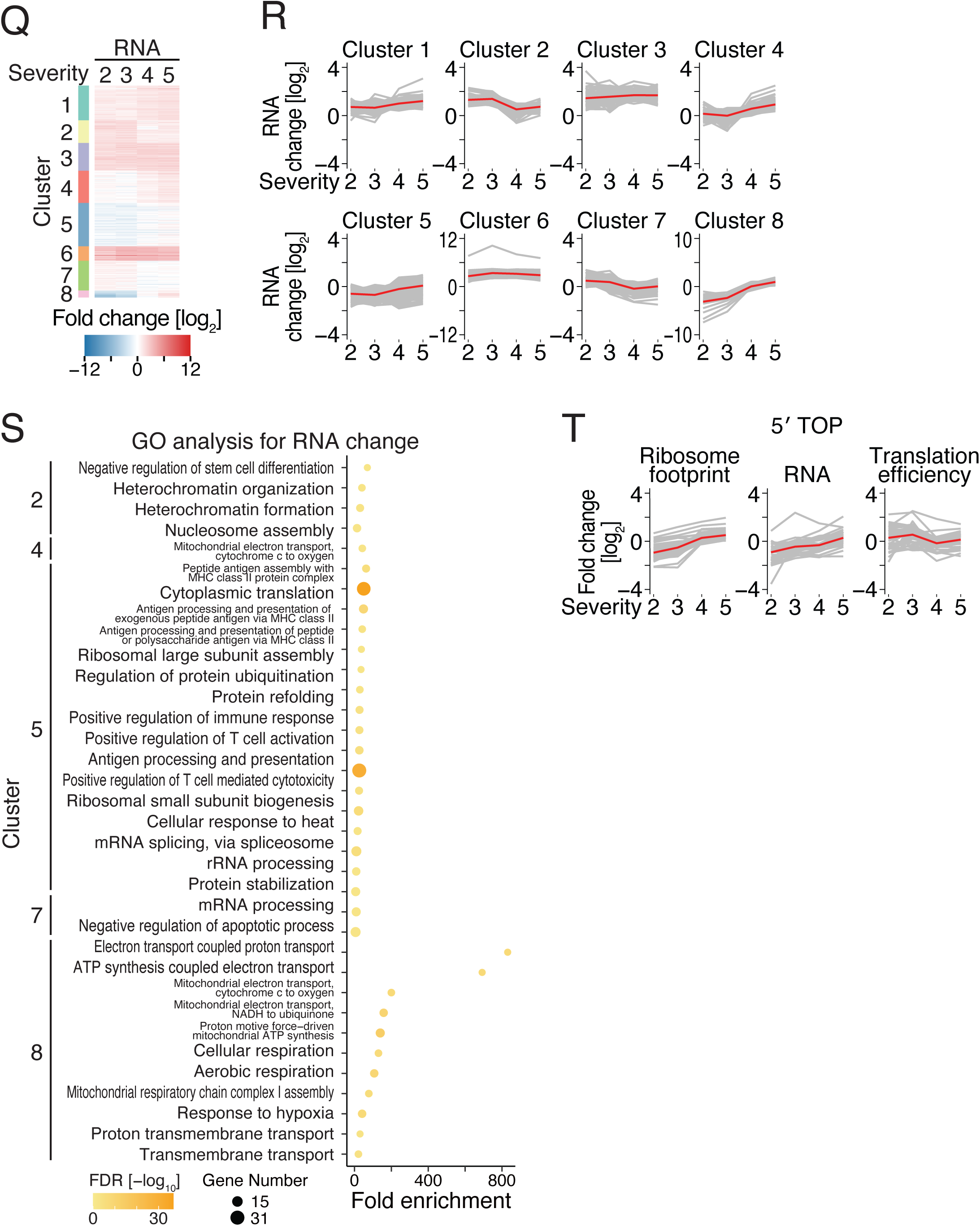
Translation landscape revealed by HT-Thor-Ribo-Seq for COVID-19-infected patients. **Related to Figure 7**. (A) The distribution of footprint length for HT-Thor-Ribo-Seq of COVID-19-infected samples. The average across 96 samples is shown as a black line. The shaded area represents ±1 standard deviation (SD). (B) Metagene plots of the 5′ end position of ribosome footprints around start (top) and stop (bottom) codons in the indicated experiments. The average across 96 samples was shown. X axis, the position relative to the start codon (0 as the first nucleotide of the start or stop codon); Y axis, footprint length; color scale, read abundance. RPM, reads per million mapped reads. (C) Pearson’s correlation coefficient (*r*) of ORF-wise counts among the non-infected samples in HT-Thor-Ribo-Seq and RNA-Seq. The *r* value scales are shown at the color bars. (D and F) Plate view of detected gene numbers (1 read or more) (D) and periodicity scores (F) for 96 samples. Color scales, indicated values. (E and G) Histogram of detected gene numbers (1 read or more) (E) and periodicity scores (G) for 96 samples. Samples with detected gene number > 3800 (dashed line in E) and periodicity score > 0.1 (dashed line in G) were used for downstream analyses. (H) Ribosome footprint, RNA, and translation efficiency changes of the indicated genes in 73 samples that passed the quality check threshold. (I) Heatmap of translation efficiency changes compared to non-infected samples. The color scale indicates the degree of fold change. The gene set detected in the translation efficiency analysis is shown. (J) Heatmap of translation efficiency changes of each severity group compared to non-infected samples. The color scale indicates the degree of fold change. The result of partitioning around medoids (PAM) clustering is shown. (K) Translation efficiency changes in each cluster (defined in J) for each severity group. The mean fold change for each group is depicted as a red line. (L) GO analysis of biological processes for the indicated clusters defined in changes of translation efficiency. Terms with FDR < 0.01 are shown. The color of the dots indicates –log_10_(FDR). (M and N) Translation efficiency changes of genes encoding OXPHOS complex subunits in the nuclear genome (M) and genes encoding proteins localized to mitochondria defined in MitoCarta3.0 (N) for each severity group. The mean fold change for each cluster is depicted as a red line. (O) Heatmap of codon-wise ribosome occupancy changes on A site (top), P site (middle), and E site (bottom) compared to non-infected samples. The mean of each severity is shown on the right. The color scale indicates the degree of fold change. (P) Ratio of ribosome footprints on the 5′ UTR and 3′ UTR. The *p* values were calculated by the two-sided Tukey-Kramer multiple comparison test. Box plots show the median (centerline), upper/lower quartiles (box limits), and 1.5× interquartile range (whiskers). (Q) Heatmap of RNA changes of each severity group compared to non-infected samples. The color scale indicates the degree of fold change. The result of partitioning around medoids (PAM) clustering is shown. (R) RNA changes in each cluster (defined in Q) for each severity group. The mean fold change for each group is depicted as a red line. (S) GO analysis of biological processes for the indicated clusters defined in changes of RNA. Terms with FDR < 0.01 are shown. The color of the dots indicates –log_10_(FDR). (T) Translation efficiency, ribosome footprint, and RNA changes of mRNAs with a 5′ TOP motif for each severity group. The mean fold change for each cluster is depicted as a red line.

**Table S1. Sample information of 192 siRNA knockdown. Related to Figures 2-3.**

**Table S2. Uniprot-based annotation for 179 knocked-down genes.** Related to Figure 3.

**Table S3. Sample information of COVID-19-infected patients.** Related to Figure 7.

**Table S4. Translation efficiency and RNA changes of each severity group compared to non-infected samples.** Related to Figure 7.

**Table S5. List of the first linker used in HT-Thor-Ribo/Disome-Seq.** Related to Figures 2-7.

**Table S6. List of the A-site offsets.** Related to Figures 1, 2, 5, and 7.

